# Large-scale mouse mutagenesis identifies novel genes affecting vertebral development

**DOI:** 10.1101/2024.09.12.612224

**Authors:** Ximena Ibarra-Soria, Elizabeth Webb, John F. Mulley

## Abstract

We analyzed the International Mouse Phenotyping Consortium (IMPC) release 19 set of 8,539 phenotyped whole-gene knockouts to identify 204 genes that alter vertebral development. These genes are broadly grouped into six categories based on their phenotype: “vertebral number” (22 genes); “vertebral processes” (35 genes); “spine shape” (16 genes); “tail morphology” (73 genes); “vertebral form” (62 genes); and “somitogenesis” (24 genes), with minimal overlap between groups. Gene expression analysis of somite trios across six developmental stages show that 182 of these genes are expressed in somites, and 60% of them show variable expression during somite maturation. A further 54% show expression changes between developmental stages. Fourteen of the 204 genes affecting vertebral development have a vertebral phenotype as their only phenotype, and for 34 genes vertebral phenotypes represent ≥50% of their total phenotypes. We find no evidence for a previous association of the majority of these genes with vertebral defects, and have therefore identified an extensive set of novel candidate genes for association with vertebral malformations in humans, including vertebral fusions, numerical variation, and scoliosis.

## Introduction

Our vertebral column defines us as vertebrates, and performs dual roles in supporting the body and providing a conduit for the nervous system. Vertebrae are not uniform, and in mammals are broadly classified into five major groups: cervical, thoracic, lumbar, sacral, and caudal. These vertebrae form from bilaterally paired blocks of paraxial mesoderm called somites (Figure 1a), which bud off from the anterior of the presomitic mesoderm starting from around 3 weeks of development in humans (Carnegie stage 9), and after around 8 days post coitus in the mouse (Theiler stage 12), at a rate of one roughly every 4-6 hours in humans, and every 2 hours in mice (Tam, 1981; William et al., 2007). Somites consist of rostral and caudal portions, and each vertebra is formed from the caudal portion of one somite and the rostral portion of the following somite through a program of resegmentation (Figure 1b) (Aoyama and Asamoto, 2000; Criswell and Gillis, 2020; Remak, 1851; Ward et al., 2017). Somites are further compartmentalised into the sclerotome, which gives rise to vertebral and rib cartilage and the associated tendons and ligaments, and the dermomyotome, which will give rise to the muscles and dermis of the back (Christ et al., 2004; Draga and Scaal, 2024; Scaal, 2016). Cells within the sclerotome behave differently depending on location, and contribute to different parts of the vertebra: those in the central sclerotome differentiate in place (i.e. without migration towards the neural tube or notochord) and form the transverse process, proximal parts of the rib, and the pedicle of the vertebral arch; cells in the ventral sclerotome migrate towards the notochord and give rise to the vertebral body; dorsal sclerotome cells migrate towards the dorsal side of the neural tube and form the vertebral arches and associated spinous processes; and lateral sclerotome cells form the distal parts of the ribs (Figure 1c) (Draga and Scaal, 2024). In the mouse, vertebral ossification begins around embryonic day 14 (e14.5). The arches begin to ossify before the centra, with the process starting from two ossification centres, one in the cervical region and one in the lower thoracic region (Hautier et al., 2014), and progressing in cranial and caudal directions from these. Developing vertebrae possess primary ossification centres in both the arches and the centrum, and so each vertebra has three ossification centres (Kaplan et al., 2005).

**Figure 1.**
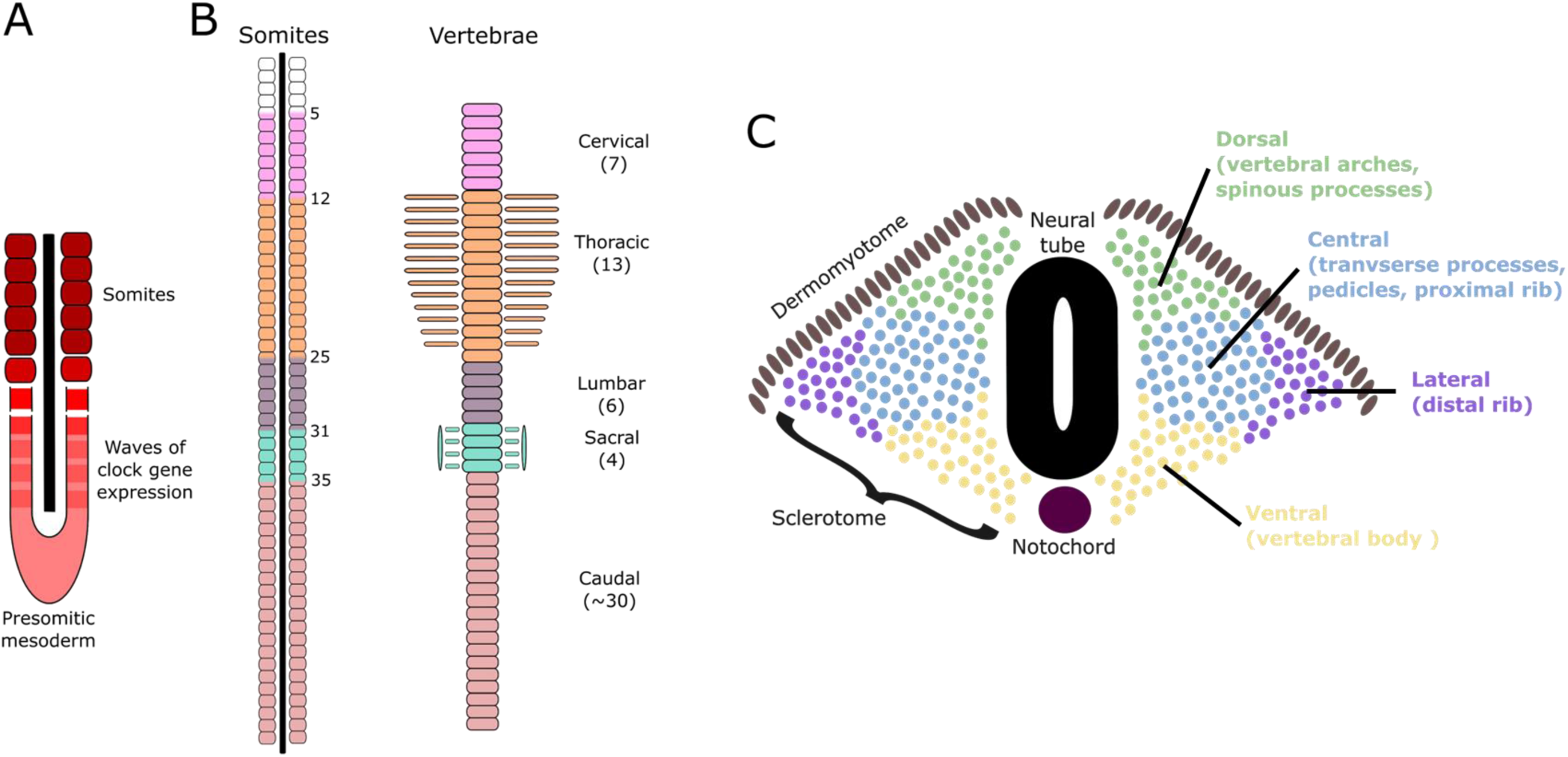
(A) Somites are blocks of paraxial mesoderm that are formed in a clock-like fashion. (B) there are seven cervical vertebrae in the mouse (formed from somites 5 to 12); 13 thoracic vertebrae (somites 12-25); 6 lumbar (somites 25-31); 4 sacral (somites 31-35) and around 30 caudal/tail vertebrae (somites 35 onwards). Vertebral boundaries are in the middle of somites, as each vertebra is formed from the rostral portion of one somite and the caudal portion of the preceding somite through a program of resegmentation. (C) Within somites, the dermomyotome gives rise to the back muscles and dermis, and the sclerotome, comprising dorsal, ventral, central and lateral portions, gives rise to the spinous processes; vertebral bodies; transverse processes, proximal ribs and pedicles; and distal ribs.

Each vertebra therefore requires the proper alignment and interaction of two opposing somites. Given the complexity of the processes of somitogenesis, resegmentation, sclerotome differentiation, and vertebral ossification, it is not surprising that things can sometimes go wrong, resulting in major health impacts. Malformations of the spine can be classified into three major groups: (i) neural tube defects, where the neural tube fails to close; (ii) defects or failures of formation, where a structural element of one or more vertebrae is missing (resulting in hemivertebrae), or where chondrification or ossification centres do not form properly (resulting in wedge vertebrae); and (iii) segmentation defects, where two or more adjacent somites fail to separate or resegment, resulting in block or bar vertebrae. (Kaplan et al., 2005; Trenga et al., 2016). Both formation and segmentation defects can lead to issues with spinal curvature such as congenital scoliosis (Pahys and Guille, 2018), and congenital vertebral malformations are thought to occur at a rate of around 0.13-0.5 per 1,000 births or more (Brand, 2008; Giampietro et al., 2009; Szoszkiewicz et al., 2024), with hemivertebrae at 1-10 per 10,000 live births (Szoszkiewicz et al., 2024), and congenital scoliosis at 0.5-1 per 1,000 births (Sebaaly et al., 2022). In addition to these structural problems, there can also be deviations from the “typical” number of vertebrae within a species through meristic changes or homeotic transformation from one vertebral type into another. Where this transformation is not complete, transitional vertebrae bearing characteristics of two adjacent vertebral types (e.g. thoracic/lumbar, lumbar/sacral) are formed. Numerical variation has been suggested to occur in between 7.7-20% of human patients (Akbar et al., 2010; Hahn et al., 1992; Tins and Balain, 2016), and transitional vertebrae may occur at anywhere between 1 and 30% of patients (Akbar et al., 2010; Chang and Nakagawa, 2004; Delport et al., 2006; McCulloch and Waddell, 1980; O’Driscoll et al., 1996; Tureli et al., 2014).

The mouse has long been a model of vertebral development. It has been almost one hundred years since Nadine Dobrovolskaya-Zavadskaya described the *short-tail* phenotype in mice heterozygous for a mutation in the *Brachyury* gene (Korzh, 2023; Korzh and Grunwald, 2001), and over 70 years since a series of research projects quantified the extent of numerical variation in the mouse spine (Green, 1953, 1951; Green and Russell, 1951; Grüneberg, 1954, 1950; McLaren and Michie, 1956, 1955, 1954) and demonstrated that the uterine environment could alter vertebral patterning (McLaren and Michie, 1958a, 1958b). Mouse mutants provided support for the role of Hox genes in vertebral patterning (Burke et al., 1995; Carapuço et al., 2005; Krumlauf, 1994; Wellik, 2007; Wellik and Capecchi, 2003), and mouse experiments highlighted the potential for interruption of this process by *in utero* exposure to retinoic acid (Kessel and Gruss, 1991). Despite this long history of research, there is still a great deal we can learn about mammalian vertebral patterning from the mouse.

The International Mouse Phenotyping Consortium (IMPC, https://www.mousephenotype.org/) is a multinational effort to produce and phenotype whole gene knock out mouse lines, with the eventual aim of generating a null mutation for every mouse gene (Brown et al., 2018; Brown and Moore, 2012; Groza et al., 2023). The mice generated to date have been used to find traits showing sexual dimorphism (Karp et al., 2017), and to identify essential genes (Dickinson et al., 2016) or those associated with: hearing loss (Bowl et al., 2017); metabolic diseases (Rozman et al., 2018); bone mineral density (Swan et al., 2020); and eye development (Chee et al., 2023; Moore et al., 2018). Here we use the May 2023 IMPC release (release 19) to identify genes that result in altered vertebral development.

## Results

We analysed the May 2023 IMPC release (release 19), which comprises 8,539 phenotyped genes and nearly 60,000 phenotype hits. Given these numbers, some degree of pleiotropy is to be expected, and indeed, the average number of phenotypes per gene is 4.43 (median 3), ranging from 0 to 77, with 1,460 genes having no phenotype assigned, and 1,282 genes a single phenotype (Figure 2).

**Figure 2.**
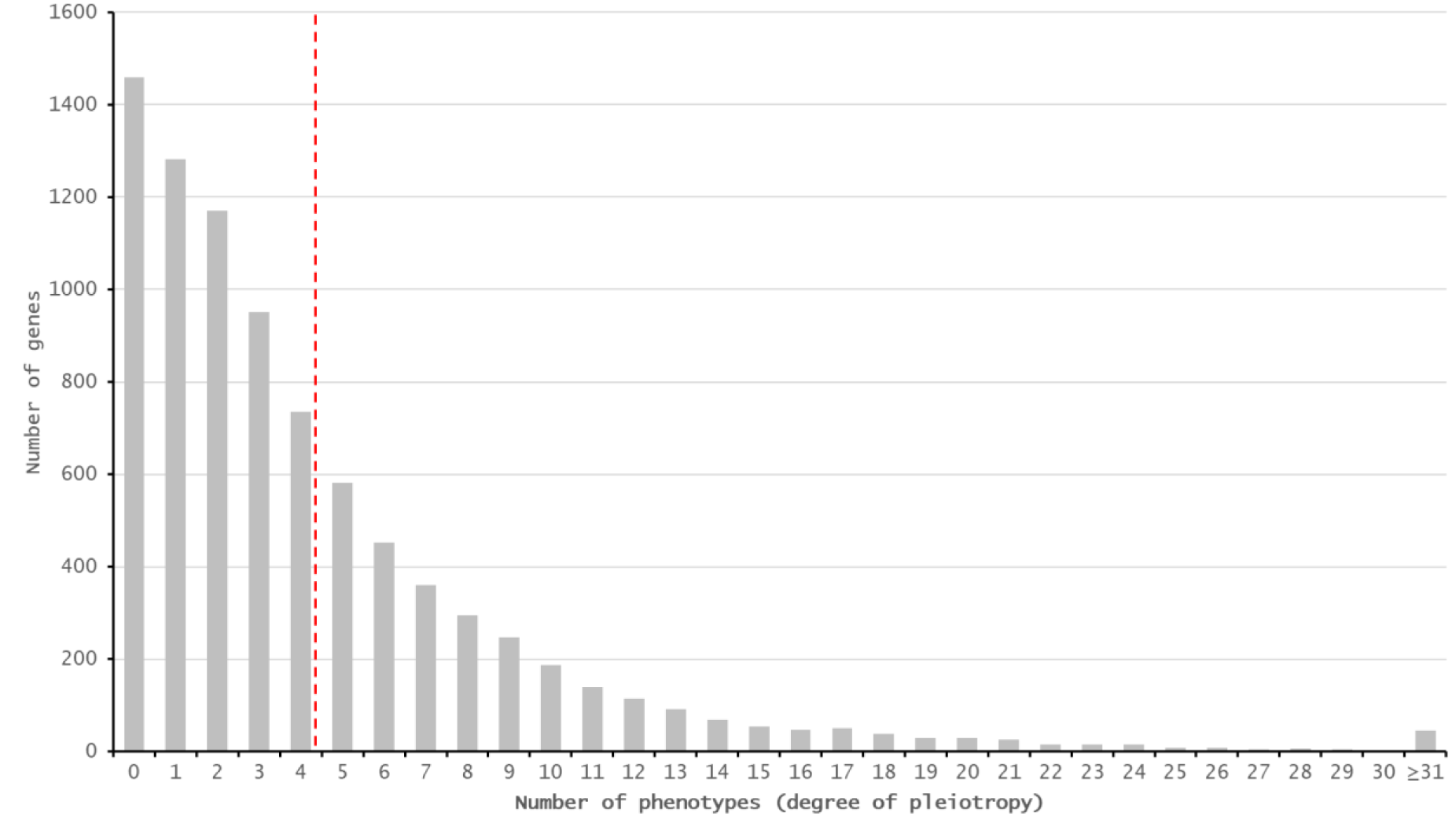
The majority of genes phenotyped by the IMPC are associated with more than one phenotype. Red line = mean (4.43). Median is 3.

Not every gene has been tested for every phenotype, and some anatomical systems are better represented than others. Of the 7,804 genes that have been tested for a skeletal phenotype, 945 showed a significant effect (Figure 3). From these, 864 showed a skeletal phenotype when homozygous, 316 when heterozygous, and 22 hemizygous (i.e. are located on the X chromosome) (Figure 3). This total is higher than 945 because some genes have an effect when both homo- and heterozygous.

**Figure 3.**
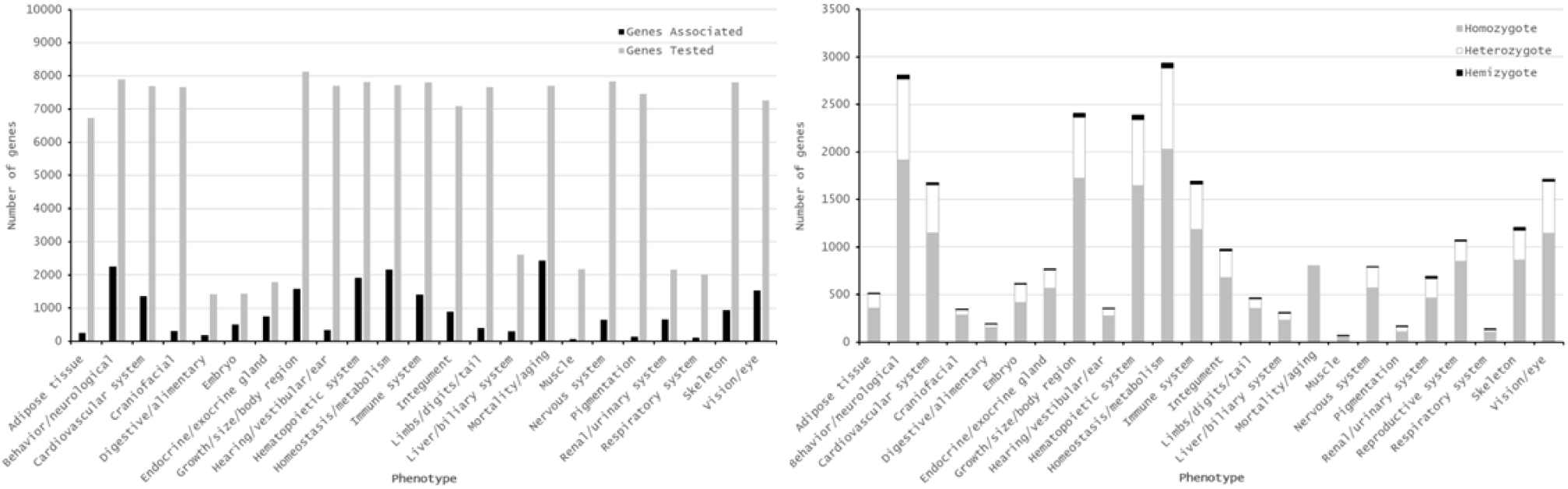
Number of genes tested and shown to be associated with higher level phenotypes (left), and zygosity of associated mutations (right). Nearly 8,000 genes have been assessed for a skeletal phenotype, and 945 show a significant effect. The majority of these genes only show a significant association in homozygous mutants.

The IMPC pipeline involves digital X-ray imaging of immobilised mice at 14 weeks of age, with results based on visual analysis of a minimum of four males and four females for 53 different IMPReSS (**I**nternational **<u>M</u>**ouse **<u>P</u>**henotyping **<u>Re</u>**source of **<u>S</u>**tandardised **<u>S</u>**creens) parameters (Groza et al., 2023). Also in week 14, body composition of a minimum of seven males and seven females is assessed using a DEXA (**<u>D</u>**ual **<u>E</u>**nergy **<u>X</u>**-ray **<u>A</u>**bsorptiometry) analyser, and body length is measured. At 9 weeks of age, mice are assessed for obvious physical, behavioural, and morphological abnormalities under the CSD (Combined SHIRPA (**<u>S</u>**mithKline, **<u>H</u>**arwell, **I**mperial College, **<u>R</u>**oyal Hospital, **<u>P</u>**henotype **<u>A</u>**ssessment (Rogers et al., 1997)) and Dysmorphology) assessments, based on a minimum of seven males and seven females. Where mutations result in lethal or subviable strains, embryonic phenotypes are assessed based on visible morphological defects at E9.5; E12.5; E14.5-15.5; and E18.5 (gross embryo morphology). From these pipelines we identified 76 parameters of relevance to skeletal development (Figure 4), with each parameter having an average of 14.3 associated genes (median 8). Thirteen parameters (*Caudal vertebrae morphology*; *Cervical vertebrae morphology*; *Fusion of ribs*; *Lumbar vertebrae morphology*; *Missing cranial rib*; *Number of cervical vertebrae*; *Number of thoracic vertebrae* ; *Number of ribs (right); Number of ribs (left)*; *Pelvic vertebrae morphology*; *Rib morphology*; *Scoliosis*; *Thoracic vertebrae morphology*) had no genes associated with them and were removed from further analysis. We also removed two parameters associated with the ribcage (*Shape of ribcage*; *Shape of ribs*); eight associated with development of teeth or the skull (*Branchial arch morphology*; *Craniofacial morphology*; *Mandibles*; *Maxilla/Pre-maxilla*; *Skull shape*; *Teeth*; *Teeth presence*; *Zygomatic bone*); and twenty-six parameters associated with the appendicular skeleton (*Brachydactyly*; *Clavicle*; *Digit integrity*; *Femu*r; *Fibula*; *Hindlimbs – size*; *Hindpaw – shape*; *Humerus*; *Joint*s; *Limb Bud Morphology*; *Limb morphology*; *Limb Plate Morphology*; *Number of digits*; *Pelvis*; *Polysyndactylism*; *Radius*; *Scapulae*; *Syndactylism*; *Syndactyly*; *Tibia*; *Tibia length (long)*; *Tibia length (short)*; *Ulna*). The *Body length* parameter was also removed as this is perhaps more equivalent to a human “height” phenotype and is likely influenced by other factors beyond vertebral changes, as evidenced by the over 200 genes associated with this parameter (Figure 4). Full details of excluded parameters, including the relevant IMPC codes can be found in the Supplemental Information.

**Figure 4.**
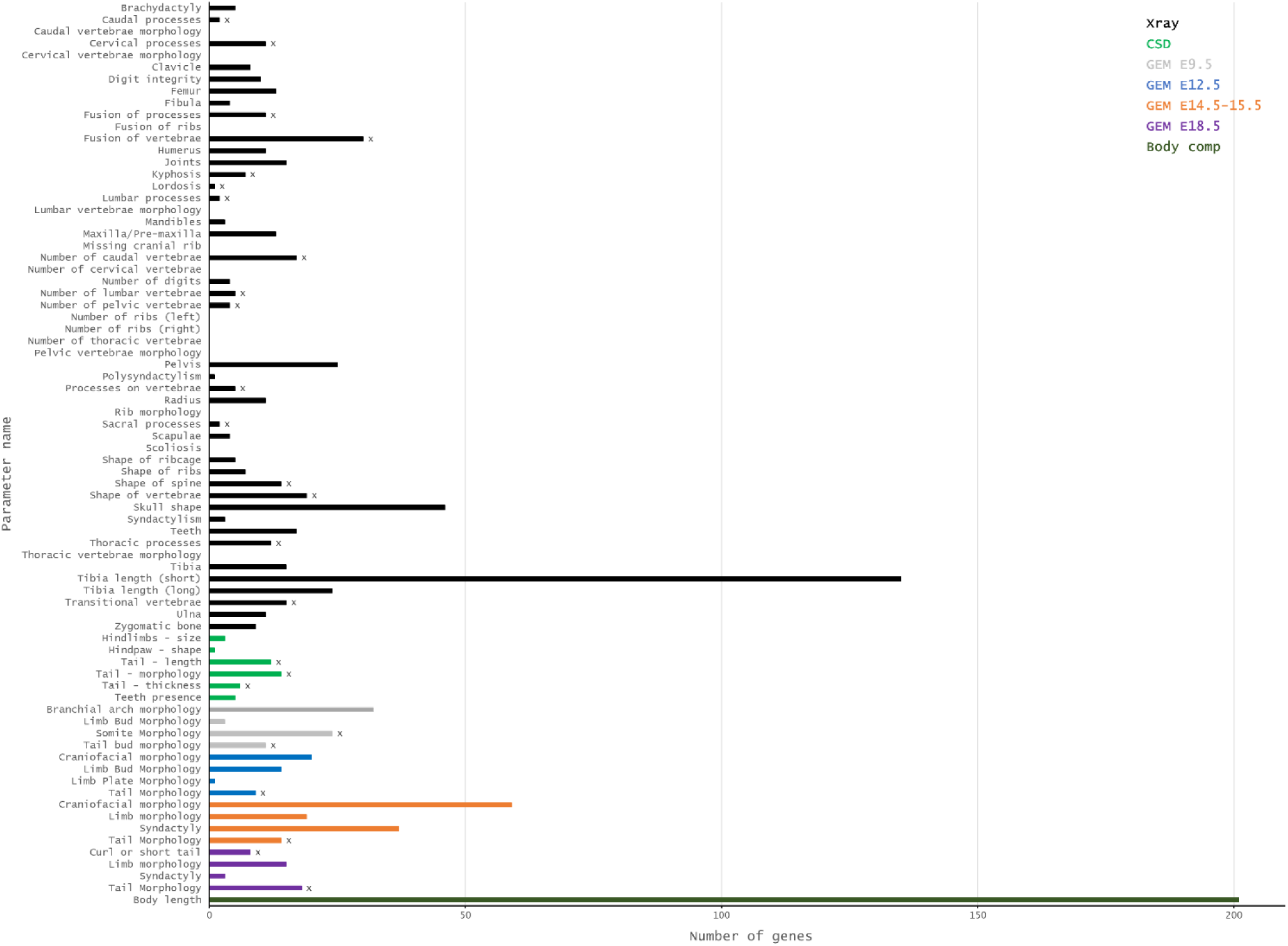
Number of genes associated with 76 skeletal parameters based on X-ray imaging of immobilised mice at 14 weeks of age (Xray); phenotype assessment and dysmorphology (CSD); gross embryo morphology at E9.5, e12.5, e14.5-15.5 and e18.5 (GEM); and body composition assessed using a DEXA (Dual Energy X-ray Absorptiometry) analyser (Body comp). Parameters marked with an x represent the 25 parameters selected for further study.

The final dataset therefore comprised 25 parameters (marked with x in Figure 4), and a non-redundant list of 204 genes (Supplemental File 1). This number represents 2.4% of genes tested in the IMPC v19 release, and 0.93% of all mouse genes, based on a total number of 21,955 genes in the GRCm39 mouse genome assembly. The number of genes associated with each of the 25 parameters varies from 1 to 30, with an average number of 10.92 genes per parameter (median 11). The majority of phenotypic changes are only apparent in homozygotes (Figure 3), and two (*Nono* and *Rlim*) are hemizygous due to their location on the X chromosome (Figure 5, Supplemental Table 1).

**Figure 5.**
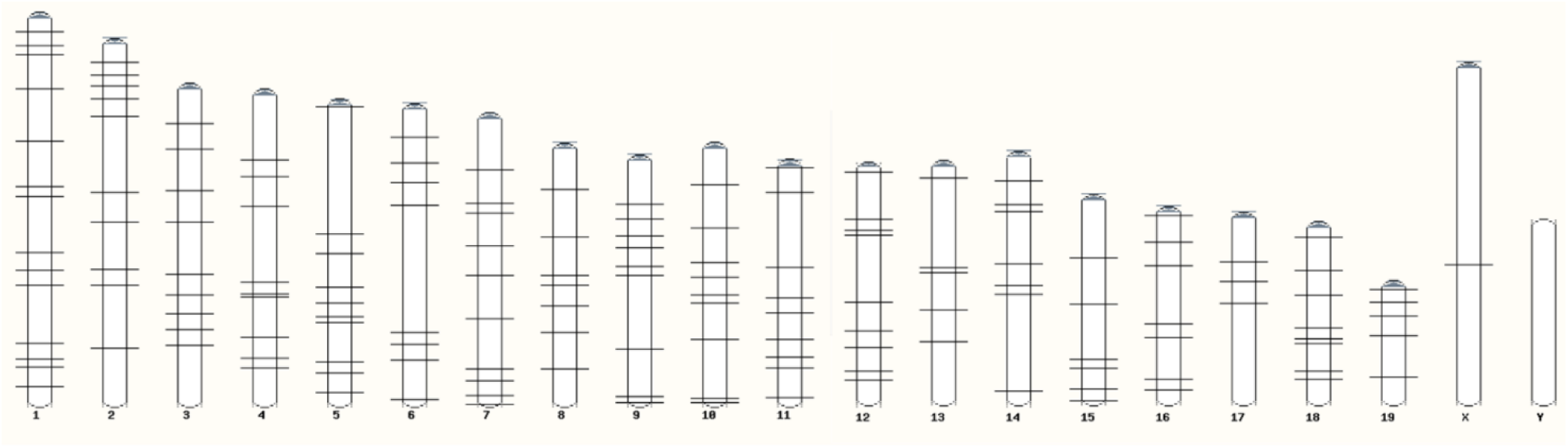
Chromosomal distribution of the 204 vertebral patterning genes from IMPC v19. Two genes (*Nono* and *Rlim*) are located on the X chromosome, but are only 2.5Mb apart and so appear as a single locus.

The 25 parameters can be assigned to six groups based on phenotype: “vertebral number” (22 genes); “vertebral processes” (35 genes); “spine shape” (16 genes); “tail morphology” (73 genes); “vertebral form” (62 genes); and “somitogenesis” (24 genes) (Table 1, Supplemental Table 2).

**Table 1.**
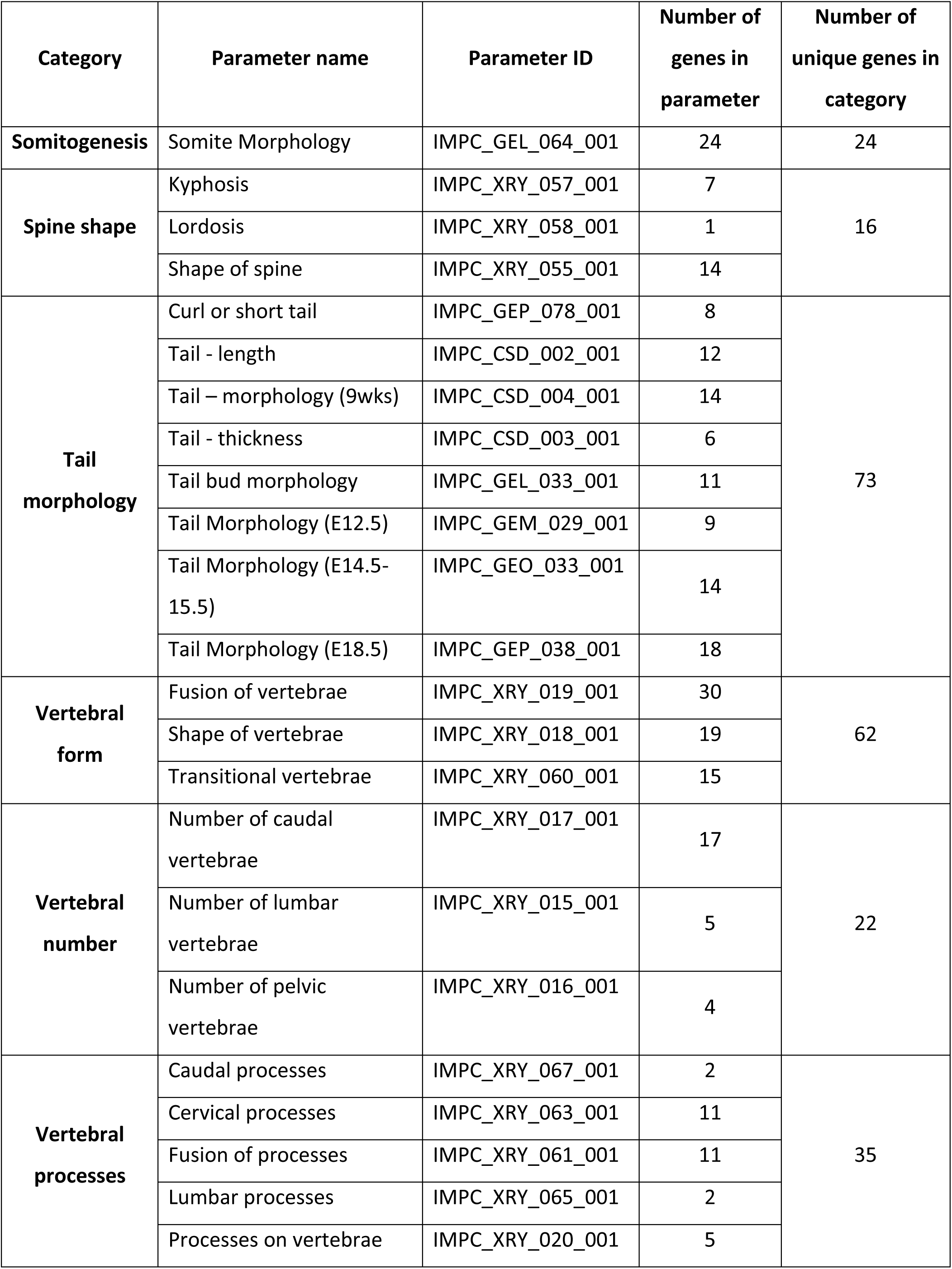
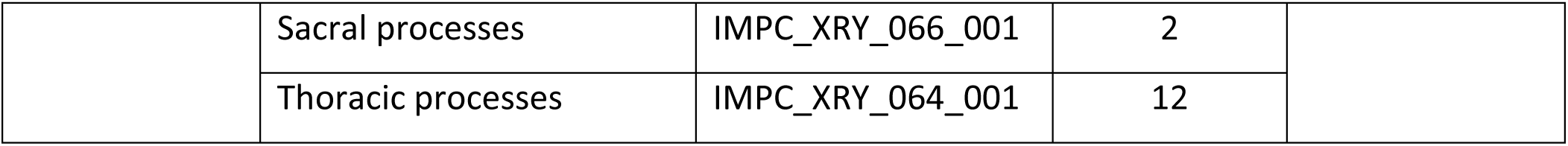
Number of genes and unique genes per IMPCR parameter and general vertebral category.

There is minimal overlap between groups (Figure 6), with the largest overlaps between “vertebral processes” and “vertebral form” (7 genes shared), and “somitogenesis” and “tail morphology” (5 genes shared). The “vertebral number” group contains the most unique genes (20/22 or 91%).

**Figure 6.**
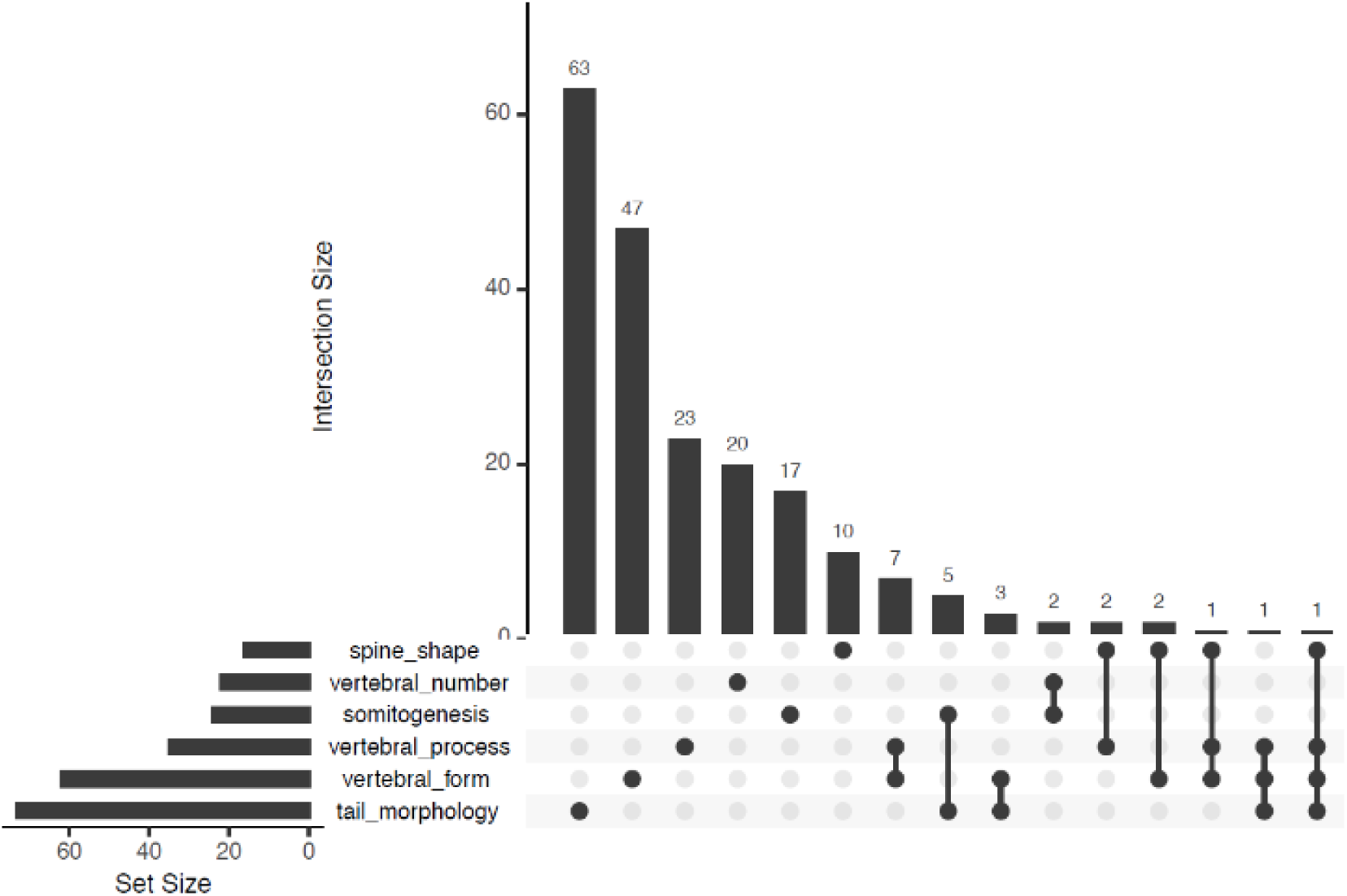
The 204 genes with a vertebral phenotype can be classified into six groups, with minimal overlap between them.

The majority of these groups show no significant enrichment for particular Gene Ontology ‘biological process’ terms, with three notable exceptions: for the ‘Somitogenesis’ group, there is enrichment for genes involved in tissue morphogenesis and neural tube development; the ‘Vertebral form’ group shows enrichment for limb morphogenesis and appendage development; and the ‘Tail morphology’ group shows enrichment for morphogenesis; skeletal system development; epidermis development; neural tube development; and pattern specification.

Additionally, the ‘Vertebral processes’ group is enriched for genes involved in enzyme binding, specifically histone deacetylase binding. Comparisons to all mouse genes are not perhaps the most appropriate, as the v19 IMPC data comprises a non-random selection of 8,539 phenotyped genes. We therefore repeated the enrichment analysis with the v19 gene set as the reference and found broadly similar results (Supplemental File 2).

Simple clustering of GO terms using ReviGO identified five clusters, which can be roughly classified as ‘development’; ‘metabolism’; ‘transport’; ‘regulation’; and ‘molecular organisation’ (Supplemental Figure S1, Supplemental File 3). Analysis of protein-protein interactions using STRING identified 21 clusters (Supplemental Figure S2) with high confidence (0.70–0.89), comprising 2-4 genes, and 12 clusters at highest confidence (0.90–1.0), comprising 2-3 genes (Supplemental Figure S3). Our dataset therefore comprises genes with a diverse set of functions, and with limited interactions between them.

### Somite expression

We next sought to determine which of our 204 candidate genes were expressed during somite formation and maturation, using RNA-Seq data from dissected somite trios at six developmental stages (8, 18, 21, 25, 27, and 35 somite stages) (Ibarra-Soria et al., 2023). Twenty-two genes were not expressed in somites at these stages, and so may alter vertebral patterning later in development, or via changes to ossification (Supplemental File 4). The remaining 182 genes were skewed to higher expression levels compared to all genes, with 74.7% expressed above the median genome-wide expression level (Figure 7A).

**Figure 7.**
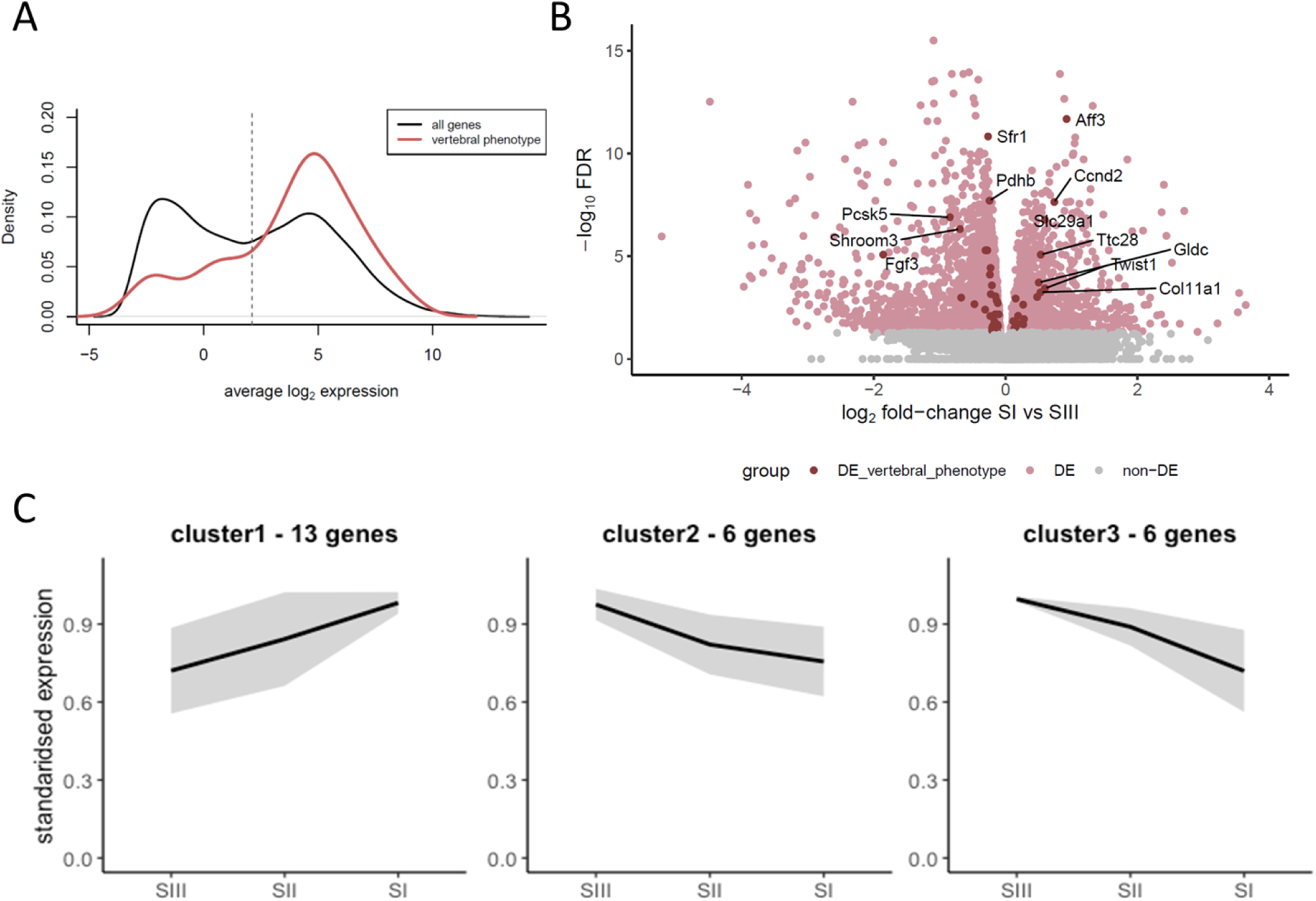
Expression of our IMPC vertebral patterning genes during somite development. 182 of 204 genes are expressed in somites, and the majority of these show elevated expression (A). 25 genes are significantly differentially expressed between somites I, II, and III, and clustering based on expression dynamics identified three patterns of expression (B, C). Roughly half of the genes (n=13) are expressed highest in the most recently segmented somite (SI) and decrease as somites mature. The other half (n=12) show the opposite behaviour.

We next explored whether these genes show variable expression across development and found nearly 60% (106/182) have a statistically significant change across somite maturation (25 genes) and/or development (98 genes). The genes with expression differences between somites I, II, and III, which might be involved in early stages of somite maturation, included equal numbers of up- and down-regulated genes, most with subtle changes (Figure 7B).

In contrast, genes that were significantly DE across developmental progression show clear expression changes between stages. This set of 98 genes can be clustered into 8 groups capturing the main patterns of expression (Figure 8). Genes in clusters 6-8, which are upregulated in somites from the lumbar and sacral skeleton, often result in tail morphology phenotypes (5/9, 4/10 and 7/11 for clusters 6-8 respectively), while vertebral form and vertebral number phenotypes involve genes from all clusters.

**Figure 8.**
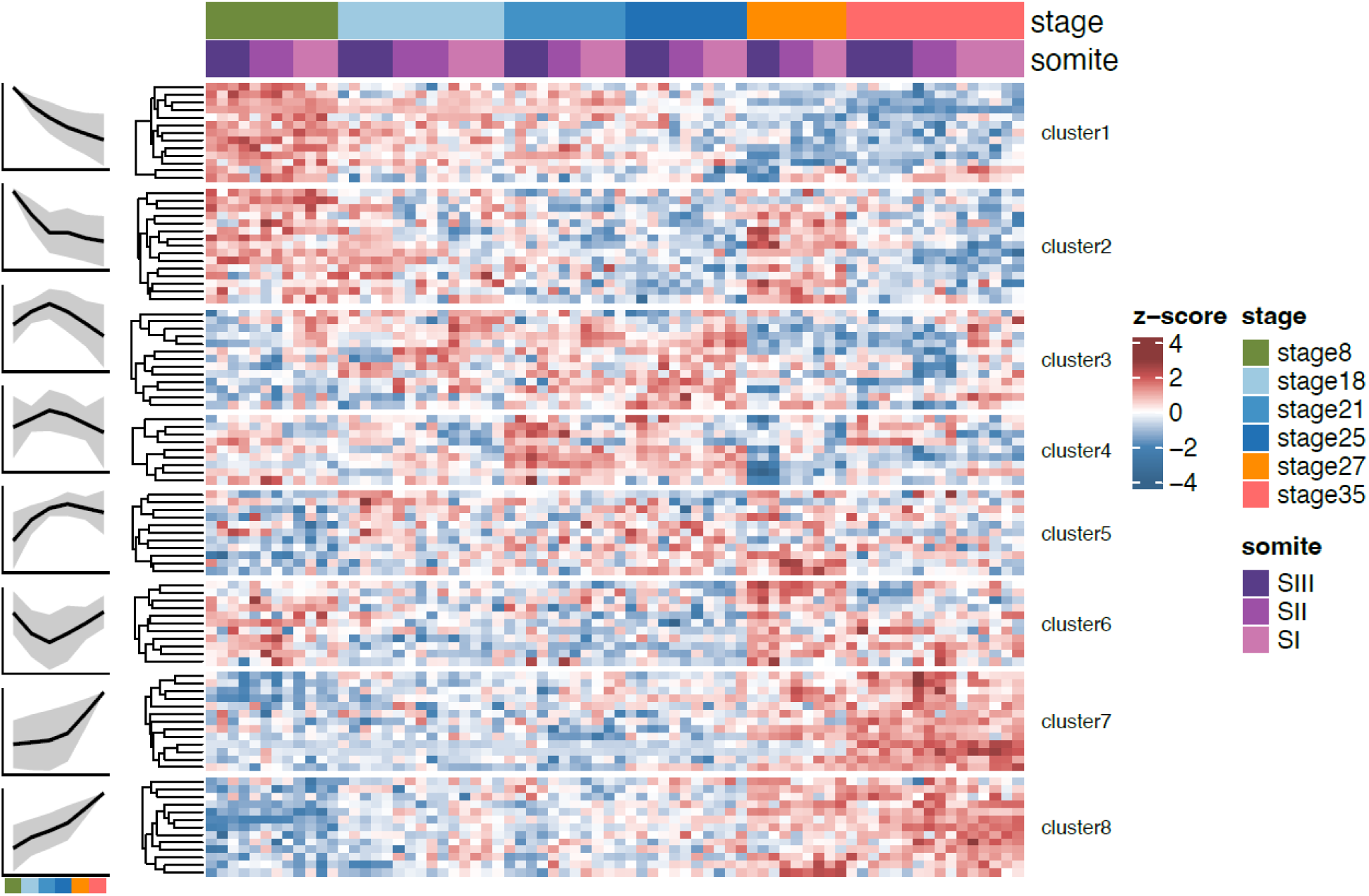
Differential expression of IMPC vertebral patterning genes during development. We find 98 genes that are differentially expressed across one or more pairs of stages of development, and cluster these into eight groups based on their expression dynamics.

### Vertebral pleiotropy

The majority of genes (n= 157) are associated with only a single vertebral parameter and so do not affect vertebral patterning more widely (Figure 9), although three genes (*Dnase1l2*, *Duoxa2*, *Fbn2*) affect five vertebral phenotypes: *Dnase1l2* is associated with ‘*Shape of spine’*; ‘*Tail – length’*; ‘*Tail* – *morphology’*; ‘*Fusion of vertebrae’*; and ‘*Fusion of processes’*; *Duoxa2* is associated with ‘*Shape of vertebrae’*; ‘*Caudal processes’*; ‘*Lumbar processes’*; ‘*Sacral processes’*; and ‘*Thoracic processes’*; *Fbn2* is associated with ‘*Tail – morphology’*; ‘*Fusion of vertebrae’*; ‘*Caudal processes’*; ‘*Lumbar processes’*; ‘*Sacral processes’* (Supplemental File 1). However, knock out of these genes results in a large number of phenotypes overall (*Dnase1l2* = 48 phenotypes; *Duoxa2* = 77 phenotypes; *Fbn2* = 43 phenotypes), far more than the 4.43 average of the entire IMPC dataset (Figure 2), and so they are disrupting embryonic development across a wide range of anatomical systems. Potentially more interesting are those genes which affect only the developing vertebral column, i.e. those where vertebral parameters represent 100% of their associated phenotypes (n=14), or those where vertebral phenotypes represent ≥50% of all phenotypes (n=34) (Figure 9, Table 2). Of this latter group, seven genes have preweaning lethality as their only non-vertebral phenotype, and all but nine are expressed in developing somites (Table 2).

**Figure 9.**
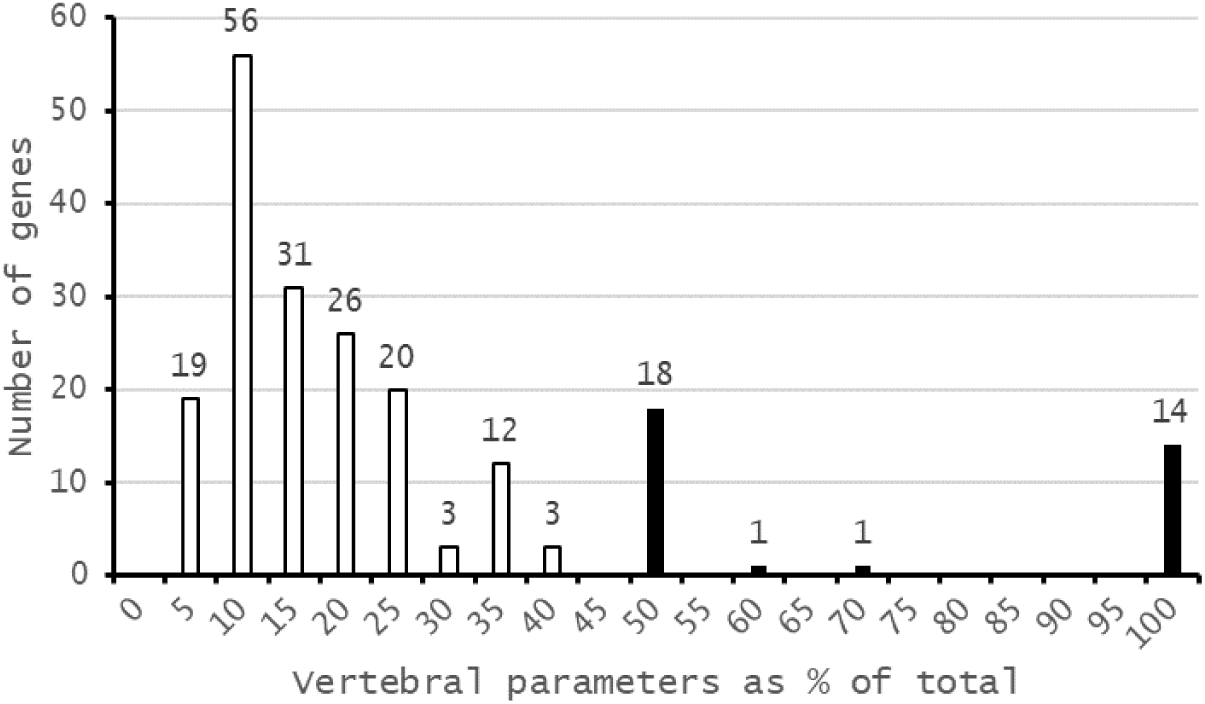
Fourteen of the 204 genes affecting vertebral development have a vertebral phenotype as their only phenotype, and for 34 genes (shown in black) vertebral phenotypes represent ≥50% of their total phenotypes. Seven of these genes have only preweaning lethality as their other phenotype.

**Table 2.** Genes where vertebral parameters are 50% or more of phenotypes (n=34), including those with only vertebral phenotypes (n=14), and those where preweaning lethality is the only other phenotype (n=7) All but nine genes are expressed in developing somites, although they do not always show differential expression (DE) between neighbouring somites or developmental stages.

|  | Gene symbol | Parameter name | Other phenotype(s) | Expressed in somites? | DE in somites | DE between stages |
| --- | --- | --- | --- | --- | --- | --- |
| Vertebral phenotype only | *1110059G10Rik* | Transitional vertebrae | - | Yes | NA | cluster2 |
|  | *4932438H23Rik* | Thoracic processes | - | No | NA | NA |
|  | *Abca4* | Fusion of vertebrae | - | Yes | NA | NA |
|  | *Fggy* | Fusion of vertebrae | - | Yes | cluster1 | cluster4 |
|  | *Gm13547* | Fusion of vertebrae | - | No | NA | NA |
|  | *H2-Eb1* | Shape of spine | - | Yes | NA | NA |
|  | *Kdm7a* | Transitional vertebrae | - | Yes | NA | cluster5 |
|  | *Kera* | Cervical processes | - | No | NA | NA |
|  | *Ppp1r42* | Fusion of vertebrae | - | No | NA | NA |
|  | *Spopl* | Thoracic processes | - | Yes | NA | NA |
|  | *Tmc6* | Fusion of vertebrae | - | Yes | cluster2 | cluster1 |
|  | *Ushbp1* | Fusion of vertebrae | - | Yes | NA | cluster2 |
|  | *Xndc1* | Fusion of processes | - | Yes | NA | NA |
|  | *Zscan2* | Fusion of vertebrae | - | Yes | NA | NA |
| **Vertebral phenotype(s)**<br>**+**<br>**preweaning lethality only** | *Chd1l* | Tail – length | Prewearing lethality, incomplete penetrance | Yes | NA | cluster5 |
|  | *Gldc* | Thoracic processes | Prewearing lethality, complete penetrance | Yes | cluster2 | cluster1 |
|  | *L3mbtl2* | Transitional vertebrae | Prewaning lethality, complete penetrance | Yes | NA | cluster1 |
|  | *Sel1l* | Fusion of vertebrae | Prewaning lethality, complete penetrance | Yes | NA | NA |
|  | *Supt5* | Transitional vertebrae | Prewaning lethality, complete penetrance | Yes | NA | cluster4 |
|  | *Vps53* | Thoracic processes | Prewaning lethality, complete penetrance | Yes | NA | NA |
|  | *Mir320* | Tail – length;<br>Tail - thickness | Prewaning lethality, complete penetrance | No | NA | NA |
| **Vertebral and non-vertebral phenotypes** | *Cbx5* | Fusion of processes;<br>Fusion of vertebrae;<br>Transitional vertebrae | Increased grip strength;<br>Decreased circulating glucose level | Yes | NA | NA |
|  | *Cdcp3* | Tail – thickness | Abnormal eye morphology | Yes |  | NA |
|  | *Dcdc2c* | Shape of vertebrae | Increased heart weight | No | NA | NA |
|  | *Hoxc12* | Tail – length | Tremors | Yes | NA | cluster2 |
|  | *Il12a* | Fusion of vertebrae | Increased circulating HDL cholesterol level | No | NA | NA |
|  | *Mbd1* | Lordosis;<br>Transitional vertebrae | Female infertility;<br>Increased mean corpuscular hemoglobin | Yes | NA | cluster5 |
|  | *Pramel17* | Tail – morphology | Enlarged heart | Yes | NA | NA |
|  | *R3hdm1* | Number of lumbar vertebrae;<br>Number of pelvic vertebrae | Small testis;<br>Abnormal testis morphology | Yes | NA | NA |
|  | *Ralb* | Thoracic processes | Increased leukocyte cell number | Yes | NA | cluster3 |
|  | *Scn3b* | Fusion of vertebrae | Abnormal bone structure | No | NA | NA |
|  | *Sh2d5* | Thoracic processes | Decreased circulating creatinine level | Yes | NA | cluster7 |
|  | *Slc6a5* | Number of lumbar vertebrae;<br>Number of pelvic vertebrae | Increased mean corpuscular hemoglobin concentration;<br>Prewaning lethality, complete penetrance | No | NA | NA |
|  | *Vgll3* | Shape of spine | Decreased startle reflex | Yes | NA | cluster6 |

Three of these genes (*Fggy, Gldc, Tmc6*) are differentially expressed during somite maturation (i.e. between somites I, II, and III), and 14 genes are differentially expressed between at least one pair of developmental stages.

### Identification of novel genes affecting vertebral development

Of the 34 genes where vertebral phenotypes represent ≥50% of all phenotypes associated with the knockout, five of them (*1110059G10Rik*, *Gm13547*, *Xndc1*, *Cdcp3*, *Pramel17*) retrieved zero results in PubMed, and another five (*4932438H23Rik*, *Ppp1r42*, *Zscan2*, *Supt5*, *Dcdc2c*) have ≤5 results. Such findings are to be expected, as the IMPC specifically focusses on phenotyping genes for which there is little to no functional annotation (the so-called “dark genome” (Lloyd et al., 2020)). The remaining genes range between 6 and 1,099 PubMed entries, with an average of 204 (median 96). Searching for the gene name with the Boolean operator “AND” and either ‘vertebra*’ or ‘somite’ did not identify any previous publications linking these genes with either somitogenesis or changes to vertebral anatomy. Searching for “gene name AND skeletal” resulted in a few relevant results for a subset of genes, and these are detailed further in the discussion section.

In early 2024, Szoszkiewicz et al. published a review of 118 genes they found to be involved in congenital vertebral malformations in humans (Szoszkiewicz et al., 2024). Of these, none overlap with our subset of 34 genes where vertebral phenotypes are ≥50% of all phenotypes, and only 10 (*AFF4, CELSR1, COL11A1, FUZ, GDF11, SHROOM3, SLC26A2, SLC29A3, SLC35D1, VANGL2*) are found in our larger set of 204 genes. Our dataset therefore comprises an extensive number of novel genes that affect vertebral patterning.

We identified 16 genes that alter spine shape, including kyphosis (*Kcnv2*); lordosis (*Mbd1*); overall spine shape (which reflects spine curvature, or scoliosis, *Ccnd2, Dnase1l2, H2-Eb1, Nisch, Pabpc4, Selenok, Uchl1, Vgll3*); or both kyphosis and overall spine shape (*Ropn1l, Sik3, Slc20a2, Tpte, Tram2, Wdr37*). These genes have between 20 and 2,498 results in PubMed (mean 355, median 106), and searches for “gene name AND kyphosis”; “gene name AND lordosis”; and “gene name

AND scoliosis” returned no results for all genes except *Uchl1*, which has previously been associated with the development of scoliosis as part of a wider range of health issues associated with a novel form of Behr syndrome (McMacken et al., 2020). These 16 spine shape genes are associated with between 1 and 5 vertebral phenotypes, and between 1 and 77 overall phenotypes. Fendri et al. previously identified 145 differentially-expressed genes (86 upregulated, 59 downregulated) in primary osteoblasts from spinal vertebrae of Adolescent Idiopathic Scoliosis (AIS) patients compared to unaffected controls (Fendri et al., 2013), and the only overlap to our dataset is the *paired-like homeodomain transcription factor 1* (*Pitx1*), which they find to be downregulated, and which we find to result in the formation of transitional vertebrae. We have therefore identified 15 novel genes that alter spine shape.

## Discussion

The knockout mice generated by the International Mouse Phenotyping Consortium offer great potential to identify novel genes underlying important human health issues. Vertebral malformations are a key part of many human syndromes, including Klippel-Feil syndrome, which affects 1 in 40,000 births, and is typically characterised by fusion of cervical vertebrae, although more caudal regions can be involved (Clarke et al., 1998; Hachem et al., 2020; Raas-Rothschild et al., 1988); the VATER/VACTERL association of vertebral defects, anal atresia, tracheoesophageal fistula with esophageal atresia, and radial or renal dysplasia (Quan and Smith, 1973), affecting 1 in 10,000 to 1 in 40,000 infants (Solomon, 2018, 2011); spondylocostal dysostoses, affecting 1 in 40,000 births (Berdon et al., 2011; Nóbrega et al., 2021); craniofacial microsomia/Goldenhar syndrome affecting between 1 in 35,000 to 1 in 56,000 births; and many others (Giampietro et al., 2009). Vertebral malformations can also present on their own, where they may be associated with congenital scoliosis, which affects 0.5 to 1 in 1,000 births (Sebaaly et al., 2022; Takeda et al., 2018), and lower back pain (Hopkins and Abbott, 2015; Jat et al., 2023; Nardo et al., 2012). Low back pain is the leading cause of disability in most countries, affecting some 619 million people in 2020, with a projection for this to rise to 843 million people by 2050 (Ferreira et al., 2023).

Our analysis of the International Mouse Phenotyping Consortium project data (release 19) has identified 204 genes that alter vertebral patterning when mutated, representing 0.93% of all mouse genes. Of these, the greatest number affect tail morphology (73 genes), suggesting that this part of the murine axial skeleton is perhaps most susceptible to (and tolerant of) change. Overall vertebral form, encompassing vertebral shape, fusion of adjacent vertebrae, and transitional vertebrae, is the next largest category with 62 genes. The majority of the 204 genes are associated with multiple phenotypes, both vertebral and non-vertebral, with a mean number of 12 phenotypes per gene (median 8), and for only 34 genes do vertebral phenotypes represent ≥50% of the assigned phenotypes. We could find no evidence of a previous association of these 34 genes with somite or vertebral development, but some did show intriguing links to relevant developmental processes. For example, the *Spopl* paralog *Spop* has been shown to be involved in skeletal development, through regulation of the hedgehog signalling pathway, specifically Indian hedgehog (*Ihh*), but a *Spopl* knockout did not produce apparent skeletal defects (Cai and Liu, 2016). However, that study was based on the Spopl^tm1(KOMP)Vlcg^ mutant, which deletes a region starting within exon 2 and ending within exon 11 between positions 23,401,253-23,435,530 of chromosome 2, whereas the IMPC phenotype is based on Spopl^tm1a(EUCOMM)Wtsi^ mutation (chr2:23,432,934-23,433,814) and deletes only a single exon (exon 5). It is possible that the vertebral arch phenotype identified in the IMPC data was too subtle to be seen in those experiments.

The microRNA *Mir320* is in the MIPF0000163 mir-320 gene family, along with human *MIR-320A*, *B*, *C* and *D*, which have been shown to be expressed in mesenchymal stem cells and which may play a role in the switch between the formation of bone marrow osteoblasts and adipocytes (Hamam et al., 2014). *Il-12* mutations decrease osteogenesis (Xu et al., 2020), and *Mbd1* is expressed in chrondrocytes (Ząbek et al., 2022). *Kdm7a* is also involved in bone development, especially adipocyte and osteoblast differentiation (Yang et al., 2019), and has been claimed to alter development of the axial skeleton in mice through alteration of the expression of posterior Hox genes (Higashijima et al., 2020; Li et al., 2023). *Cbx5* is involved in osteoblast differentiation, and may alter Hox gene expression through modification of chromatin structure (Dashti et al., 2024; van Wijnen et al., 2021), in a similar way to that of the paralogous *Cbx2* (Sato et al., 2020). Vgll3 has a role in osteogenic and myogenic differentiation (Gabay Yehezkely et al., 2020; Yuan et al., 2022) as does the related *Vgl-2* in chicken and zebrafish (Bonnet et al., 2010; Mann et al., 2007). The carbohydrate kinase *Fggy* is expressed in skeletal muscle, and has been suggested to have a role in muscle cell differentiation (Smith et al., 2021), and therefore the somite expression of this gene (Table 2) suggests a role in the dermomyotome, which gives rise to skeletal muscles. *Ralb* is also expressed in skeletal muscle (Chen et al., 2019; Wildey et al., 1993). *CHD1L* has been implicated in skeletal defects in stillborn fetuses, nervous system development, and short stature (Dou et al., 2017; Henrie et al., 2018; Workalemahu et al., 2024), and Gldc is expressed in neuromesodermal progenitors (Chang et al., 2022; Gouti et al., 2017). Finally, *Hoxc12* has been implicated in vertebral development, through disruption of the genomic region in deletion of the *Hotair* long non-coding RNA (lncRNA) (Amândio et al., 2016), although given the role of Hox genes in axial pattering (Burke et al., 1995; Carapuço et al., 2005; Krumlauf, 1994; Wellik, 2007), such a role is not surprising, and it is likely that the mildness of the phenotype (short tail) can be explained through compensation by the paralogous *Hoxd12*. We find that these short-tail *Hoxc12* knockout mice (Figure 10) have fewer caudal vertebrae than wild type controls (26.5 vs 27.8 respectively), based on an analysis of the IMPC X-ray images for 12 mutant and 15 wildtype mice.

**Figure 10.**
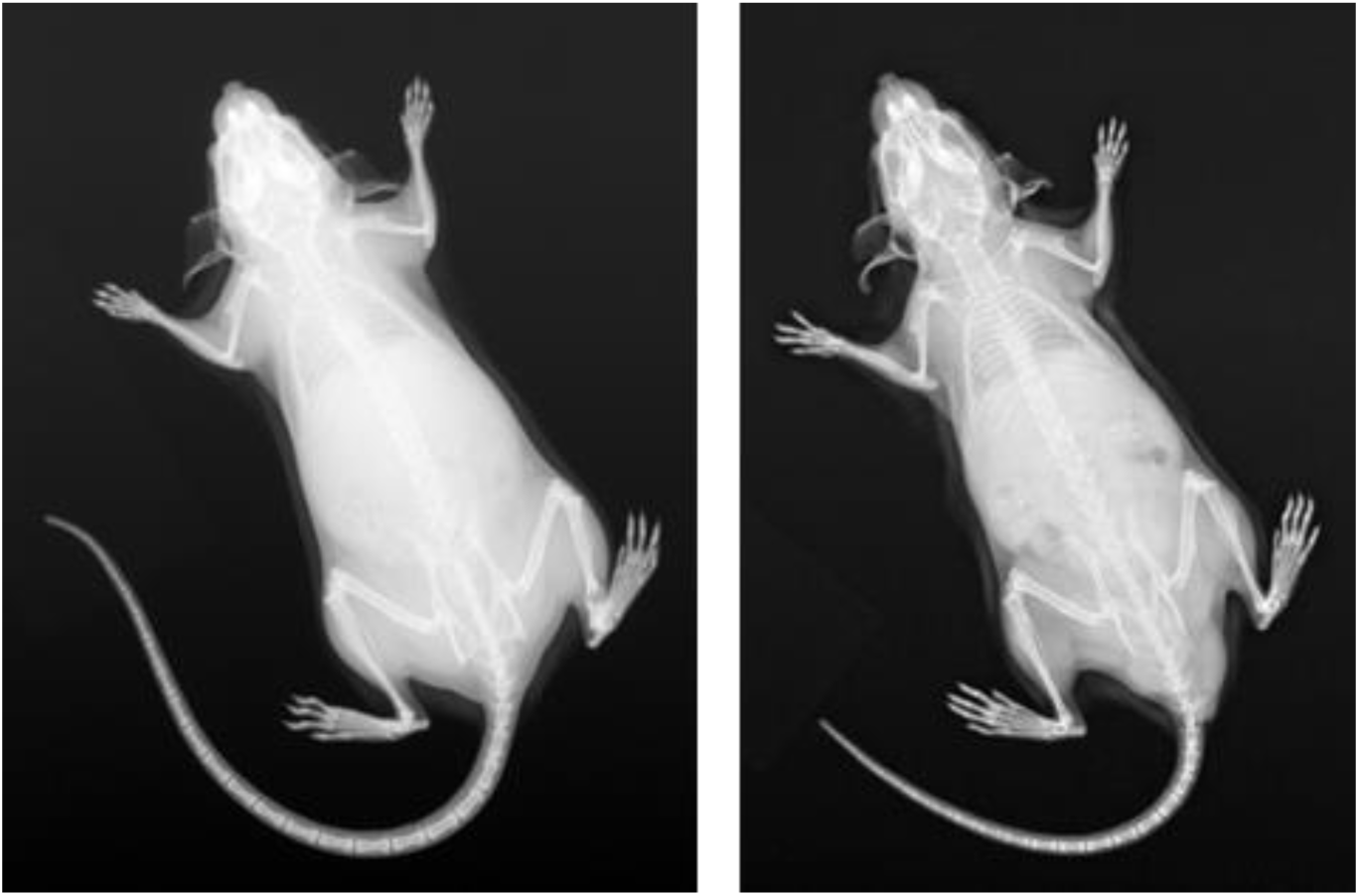
Wildtype (left) and *Hoxc12* knockout (right) mice. The *Hoxc12* knockouts have significantly shorter tails, and fewer caudal vertebrae.

Our gene ontology and network analyses suggested that the genes we have identified represent members of a diverse range of biological processes in vertebral specification, development, and ossification, the majority of which (182 of 204) are expressed in developing somites in at least one of the six developmental stages we looked at. Our dataset therefore comprises genes with a diversity of roles in vertebral development, covering somitogenesis, the specification of vertebral identity, sclerotome migration and differentiation, and later chondrogenesis and ossification.

Given the large number of knockout phenotypes that alter vertebral shape and vertebral processes (62 and 35 respectively, or 87 overall when duplicate genes found in both categories are removed), it seems likely that the sclerotome differentiation and subsequent chondrogenesis and ossification are some of the most sensitive parts of vertebral development.

Within our set of 204 genes, nine (*Barx2, Cyb561, Fbn2, Gal3st1, Hip1, Lrrk1, Micu1, Npr2, Sirt3*) have a vertebral phenotype and affect gait (MP:0001406 ‘abnormal gait’); seven genes (*Cyp27b1, Duoxa2, Fbn2, Lrrk1, Sik3, Slc29a1, Tm9sf4*) affect both vertebral development and the ribcage (MP:0000150 ‘Abnormal rib morphology’; MP:0004624 ‘abnormal thoracic cage morphology’); and eight (*Cyp27b1, Fbn2, Lrrk1, Runx2, Sik3, Ttc28, Twist1, Ube2g1*) result in abnormal development of the pelvic girdle (MP:0004508 ‘abnormal pectoral girdle bone morphology’). The IMPC knockout mice that we have identified therefore represent not only a powerful resource to improve our understanding of the processes underlying vertebral development, but also the links between these processes and the development of associated skeletal structures, with relevance for our understanding of a diversity of human diseases.

## Conclusions

Mammalian vertebral development is a complex process, and the loss of a single gene is often sufficient to produce extreme phenotypes such as fewer or fused vertebrae. We have identified 204 genes where the knockout phenotype results in changes to vertebral development, including 14 which only produce a vertebral phenotype, and a further 20 where vertebral phenotypes represent ≥50% of all recorded phenotypes. The fact that nearly 1% of all mouse genes can alter vertebral development reflects the complexity of the underlying processes, and demonstrates that novel cellular and molecular mechanisms could underlie vertebral phenotypes in a range of human health issues.

## Methods

### Identification of candidate genes

We used the IMPC programmatic data access portal application programming interface (API, https://www.mousephenotype.org/help/programmatic-data-access/) to download all records resulting from the digital X-ray imaging (XRY); gross embryo morphology at E9.5 (GEL); E12.5 (GEM); E14.5-E15.5 (GEO); E18.5 (GEP); combined SHIRPA ((**<u>S</u>**mithKline, **<u>H</u>**arwell, **I**mperial College, **<u>R</u>**oyal Hospital, **<u>P</u>**henotype **<u>A</u>**ssessment) and dysmorphology (CSD); and body composition (DXA) IMPC release 19 phenotyping pipelines. From these data we identified 76 parameters (defined by IMPC as “a measurement obtained during an experiment”) related to the skeleton, and then parsed these to only those that contained ≥1 gene and that pertained to the vertebral column (i.e removed those that pertained to teeth, the skull, limbs, or the ribcage), resulting in a set of 204 genes that alter skeletal development in heterozygous, hemizygous, or homozygous knockouts.

### Functional analysis

We first performed a rough clustering of Gene Ontology terms associated with these genes using ReviGO (Supek et al., 2011), based on gene annotations from the <u>D</u>atabase for <u>A</u>nnotation, <u>V</u>isualization and Integrated <u>D</u>iscovery (DAVID 2021) functional annotation tool (v2023q4, (Sherman et al., 2022), in the GOTERM_BP_FAT category and restricted to terms with modified Fisher exact p-values <0.05. We next analysed gene function enrichment using the PANTHER (<u>P</u>rotein <u>AN</u>notation <u>TH</u>rough <u>E</u>volutionary <u>R</u>elationship) classification system (Version 18.0 released 2023-08-01 (Mi et al., 2013) , for the *biological process (BP)*, *molecular function (MF)*, and *cellular component (CC)* categories, with comparison to both the full mouse gene set (21,983 genes in PANTHER v18.0) and the smaller IMPC v19 gene set (8,539 genes), focussing only on results with a false discovery rate (FDR) of p <0.05. Protein-protein interactions between candidate genes were predicted using the STRING (<u>S</u>earch <u>T</u>ool for <u>R</u>ecurring Instances of <u>N</u>eighbouring <u>G</u>enes) database(Szklarczyk et al., 2019), with MCL clustering (Brohée and van Helden, 2006), and restriction to high (0.70–0.89) or highest (0.90–1.0) confidence interactions.

#### Somite data methods

To assess whether the 204 genes might be involved in the development of the skeletal system, we checked their expression levels in the dataset from Ibarra-Soria et al., 2023 (batch corrected gene expression levels were downloaded from ArrayExpress E-MTAB-12511). We also cross-referenced against the set of genes reported as significantly differentially expressed between somites I, II and III, or between stages (downloaded from https://github.com/xibarrasoria/somitogenesis2022).

#### Gene analysis

For genes where a vertebral phenotype was their only significant phenotype, and those where vertebral phenotypes represented ≥50% of all phenoptyes, we looked for previous associations with vertebral development throughs searches of PubMed (www.pubmed.gov) using the gene name by itself, and the gene name with the Boolean operator “AND” and the search terms ‘vertebra*’, ‘somite’, or ‘skeletal’. We looked for previous links to scoliosis and spine shape in the same way, using the search terms “gene name AND kyphosis”; “gene name AND lordosis”; and “gene name AND scoliosis”. We also searched Google Scholar (www.google.com/scholar), using the gene name (in quotation marks to ensure exact match) and the above search terms, but this tended to result in large numbers of results due to spurious matches.

## Acknowledgments

The authors wish to thank Piia Keskivali-Bond, Sharon Cheng, Sara Wells, and James Brown for useful discussions, and for help with navigating IMPC data.

