## Supplemental information for "Large-scale mouse mutagenesis identifies novel genes affecting vertebral development"

See also:

Supplemental file 1 – 204 vertebral genes and their associated phenoptypes

Supplemental file 2 – Panther results for the IMPC gene set reference

Supplemental file 3 – ReviGO clustering

Supplemental file 4 – Somite trio RNA-Seq expression and clustering results

### Excluded skeletal parameters

**Parameters with no associated genes:**

*Caudal vertebrae morphology* [IMPC_XRY_074_001]
*Cervical vertebrae morphology* [IMPC_XRY_070_001]
*Fusion of ribs* [IMPC_XRY_011_001]
*Lumbar vertebrae morphology* [IMPC_XRY_072_001]
*Missing cranial rib* [IMPC_XRY_068_001]
*Number of cervical vertebrae* [IMPC_XRY_013_001]
*Number of thoracic vertebrae* [IMPC_XRY_014_001]
*Number of ribs (right)* [IMPC_XRY_008_001]
*Number of ribs (left)* [IMPC_XRY_009_001]
*Pelvic vertebrae morphology* [IMPC_XRY_073_001]
*Rib morphology* [IMPC_XRY_069_001]
*Scoliosis* [IMPC_XRY_056_001]
*Thoracic vertebrae morphology* [IMPC_XRY_071_001]

**Ribcage parameters:**

*Shape of ribcage* [IMPC_XRY_059_001]
*Shape of ribs* [IMPC_XRY_010_001]

**Tooth or skull parameters:**

*Branchial arch morphology* [IMPC_GEL_024_001]
*Craniofacial morphology* [IMPC_GEO_063_001 and IMPC_GEM_062_001]
*Mandibles* [IMPC_XRY_004_001]
*Maxilla/Pre-maxilla* [IMPC_XRY_003_001]
*Skull shape* [IMPC_XRY_001_001]
*Teeth* [IMPC_XRY_005_001]
*Teeth presence* [IMPC_CSD_068_001]
*Zygomatic bone* [IMPC_XRY_002_001]

**Appendicular skeleton parameters:**

*Brachydactyly* [IMPC_XRY_030_001]
*Clavicle* [IMPC_XRY_007_001]
*Digit integrity* [IMPC_XRY_031_001]
*Femur* [IMPC_XRY_024_001]
*Fibula* [IMPC_XRY_026_001]
*Hindlimbs – size* [IMPC_CSD_022_001]
*Hindpaw – shape* [IMPC_CSD_043_001]
*Humerus* [IMPC_XRY_021_001]
*Joints* [IMPC_XRY_027_001]
*Limb Bud Morphology* [IMPC_GEL_038_001 and IMPC_GEM_027_001]
*Limb morphology* [IMPC_GEP_084_001 and IMPC_GEO_065_001]
*Limb Plate Morphology* [IMPC_GEM_028_001]
*Number of digits* [IMPC_XRY_028_001]
*Pelvis* [IMPC_XRY_012_001]
*Polysyndactylism* [IMPC_XRY_062_001]
*Radius* [IMPC_XRY_022_001]
*Scapulae* [IMPC_XRY_006_001]

*Syndactylism* [IMPC_XRY_029_001]
*Syndactyly* [IMPC_GEP_082_001 and IMPC_GEO_067_001]
*Tibia* [IMPC_XRY_025_001]
*Tibia length (long)* [IMPC_XRY_033_001]
*Tibia length (short)* [IMPC_XRY_033_001]
*Ulna* [IMPC_XRY_023_001]

**Body length**

*body length* [IMPC_DXA_006_001]

Supplemental table 1. Chromosomal locations of genes affecting vertebral phenotypes based on mouse GRCm39

| Ensembl ID | Chromosome | Genomic location (strand) |
|---|---|---|
| *1110059G10Rik* | 9 | 122774154-122780065(-1) |
| *4932438H23Rik* | 16 | 90850823-90892010(-1) |
| *Abca4* | 3 | 121838092-121973772(1) |
| *Adamts1* | 16 | 85590715-85600001(-1) |
| *Aebp1* | 11 | 5811947-5822088(1) |
| *Aff3* | 1 | 38216407-38704036(-1) |
| *Aff4* | 11 | 53241660-53312657(1) |
| *Alg10b* | 15 | 90108514-90117674(1) |
| *Ambra1* | 2 | 91560479-91749194(1) |
| *Ankmy2* | 12 | 36207113-36247290(1) |
| *Arpc1b* | 5 | 145051025-145067515(1) |
| *Asphd1* | 7 | 126544739-126548754(-1) |
| *Atp1b1* | 1 | 164264678-164285924(-1) |
| *Atp1b3* | 9 | 96214708-96246495(-1) |
| *Barx2* | 9 | 31757340-31824758(-1) |
| *Bptf* | 11 | 106923907-107022953(-1) |
| *Brd2* | 17 | 34330997-34341608(-1) |
| *C9orf72* | 4 | 35191285-35226175(-1) |
| *Cacna1c* | 6 | 118564201-119173851(-1) |
| *Cant1* | 11 | 118297115-118309912(-1) |
| *Capza1* | 3 | 104730095-104771821(-1) |
| *Cbx5* | 15 | 103099971-103148243(-1) |
| *Ccnd2* | 6 | 127102125-127129156(-1) |
| *Cdcp3* | 7 | 130776131-130908180(1) |
| *Celsr1* | 15 | 85783130-85918404(-1) |
| *Cfd* | 10 | 79726687-79728489(1) |
| *Cgn* | 3 | 94667376-94693826(-1) |
| *Chd1l* | 3 | 97468058-97517519(-1) |
| *Chrna2* | 14 | 66372488-66390397(1) |
| *Cog6* | 3 | 52889296-52924658(-1) |
| *Col11a1* | 3 | 113824189-114014367(1) |
| *Colec10* | 15 | 54274170-54329754(1) |
| *Coq6* | 12 | 84408431-84420570(1) |
| *Cpt2* | 4 | 107761178-107780807(-1) |
| *Ctbp2* | 7 | 132589292-132726083(-1) |
| *Cx3cr1* | 9 | 119730682-119898945(-1) |
| *Cyb561* | 11 | 105824528-105844162(-1) |
| *Cyp27b1* | 10 | 126884119-126888875(1) |
| *Cysltr2* | 14 | 73263043-73286554(-1) |
| *Dbn1* | 13 | 55621242-55635924(-1) |
| *Dcbld2* | 16 | 58228806-58290090(1) |
| *Dcdc2c* | 12 | 28487794-28602398(-1) |

| | | |
|---|---|---|
| *Dis3l2* | 1 | 86631530-86977817(1) |
| *Dnase1l2* | 17 | 24659055-24662079(-1) |
| *Dpf2* | 19 | 5946544-5963038(-1) |
| *Dph6* | 2 | 114346897-114485445(-1) |
| *Duoxa2* | 2 | 122129381-122133366(1) |
| *Eng* | 2 | 32536607-32572681(1) |
| *Fbn2* | 18 | 58141695-58343559(-1) |
| *Fbrsl1* | 5 | 110509620-110596468(-1) |
| *Fbxo22* | 9 | 55116209-55131717(1) |
| *Fgf3* | 7 | 144391820-144398173(1) |
| *Fgf7* | 2 | 125876578-125933105(1) |
| *Fgfr1op2* | 6 | 146478701-146500696(1) |
| *Fggy* | 4 | 95445744-95815176(1) |
| *Flnc* | 6 | 29433255-29461882(1) |
| *Fuz* | 7 | 44545503-44552055(1) |
| *Gal3st1* | 11 | 3933636-3949326(1) |
| *Gdf11* | 10 | 128718164-128727587(-1) |
| *Gldc* | 19 | 30075847-30152829(-1) |
| *Glg1* | 8 | 111881053-111985848(-1) |
| *Glmn* | 5 | 107696833-107745754(-1) |
| *Gm13547* | 2 | 29649324-29654361(1) |
| *Gna11* | 10 | 81364558-81381024(-1) |
| *Gramd1b* | 9 | 40204529-40442679(-1) |
| *Gramd2* | 9 | 59587427-59626157(1) |
| *Gyg* | 3 | 20176248-20209481(-1) |
| *H2-Eb1* | 17 | 34524841-34535648(1) |
| *Hbs1l* | 10 | 21171878-21244797(1) |
| *Helq* | 5 | 100910011-100946464(-1) |
| *Hip1* | 5 | 135435385-135573974(-1) |
| *Hoxc12* | 15 | 102845192-102847044(1) |
| *Htr1f* | 16 | 64745092-64926217(-1) |
| *Ifi27l2a* | 12 | 103408426-103409939(-1) |
| *Il12a* | 3 | 68597977-68605880(1) |
| *Il2ra* | 2 | 11647618-11698004(1) |
| *Ino80c* | 18 | 24237814-24255010(-1) |
| *Ipo9* | 1 | 135310050-135358237(-1) |
| *Isyna1* | 8 | 71047023-71049940(1) |
| *Jmjd1c* | 10 | 66931904-67092105(1) |
| *Kcnv2* | 19 | 27299988-27314579(1) |
| *Kdm7a* | 6 | 39113557-39183723(-1) |
| *Kera* | 10 | 97442741-97449554(1) |
| *Klf7* | 1 | 64068606-64161441(-1) |
| *Krit1* | 5 | 3853184-3895564(1) |
| *L3mbtl2* | 15 | 81548090-81572516(1) |
| *Lox* | 18 | 52649139-52662939(-1) |
| *Lrrk1* | 7 | 65876660-66038098(-1) |

| | | |
|---|---|---|
| *Lum* | 10 | 97400990-97408565(1) |
| *Lyplal1* | 1 | 185819928-185849507(-1) |
| *Mau2* | 8 | 70468773-70495384(-1) |
| *Mbd1* | 18 | 74400676-74415803(1) |
| *Med11* | 11 | 70342745-70344553(1) |
| *Micu1* | 10 | 59538299-59699954(1) |
| *Mir320* | 14 | 70680950-70681031(1) |
| *Mllt10* | 2 | 18060048-18217199(1) |
| *Mmp11* | 10 | 75759056-75772330(-1) |
| *Mtch2* | 2 | 90677499-90697154(1) |
| *Ndel1* | 11 | 68712260-68762684(-1) |
| *Ndufb5* | 3 | 32791139-32805715(1) |
| *Nemf* | 12 | 69357296-69403939(-1) |
| *Nisch* | 14 | 30892887-30938903(-1) |
| *Nog* | 11 | 89191464-89193158(-1) |
| *Nono* | X | 100472924-100492197(1) |
| *Notch1* | 2 | 26347915-26406675(-1) |
| *Npr2* | 4 | 43631935-43651244(1) |
| *Nr6a1* | 2 | 38613382-38817700(-1) |
| *Opa1* | 16 | 29398152-29473702(1) |
| *Os9* | 10 | 126931519-126957000(-1) |
| *Pabpc4* | 4 | 123156144-123192718(1) |
| *Pals2* | 6 | 50087221-50175919(1) |
| *Parp4* | 14 | 56813076-56897251(1) |
| *Pcgf2* | 11 | 97579649-97591323(-1) |
| *Pcgf3* | 5 | 108609098-108654842(1) |
| *Pcsk5* | 19 | 17409683-17814996(-1) |
| *Pde5a* | 3 | 122522596-122653023(1) |
| *Pdhb* | 14 | 14296748-14303777(1) |
| *Pdss2* | 10 | 43097482-43340878(1) |
| *Pfdn1* | 18 | 36536729-36587577(-1) |
| *Pfdn5* | 15 | 102234551-102240308(1) |
| *Pfn4* | 12 | 4819022-4828813(1) |
| *Pigq* | 17 | 26145395-26163910(-1) |
| *Pitx1* | 13 | 55972864-55984005(-1) |
| *Pld5* | 1 | 175789872-176102878(-1) |
| *Plekhm1* | 11 | 103255101-103303513(-1) |
| *Ppp1r35* | 5 | 137777111-137778372(1) |
| *Ppp1r42* | 1 | 10038849-10079361(-1) |
| *Pramel17* | 4 | 101692166-101701220(-1) |
| *Psen1* | 12 | 83734926-83781973(1) |
| *Pstpip2* | 18 | 77877614-77970579(1) |
| *R3hdm1* | 1 | 128031038-128165473(1) |
| *Rad9a* | 19 | 4245195-4251661(-1) |
| *Ralb* | 1 | 119398035-119432524(-1) |
| *Rbm22* | 18 | 60693808-60705882(1) |

| | | |
|---|---|---|
| *Rbm45* | 2 | 76200328-76214112(1) |
| *Rexo1* | 10 | 80376756-80397394(-1) |
| *Rlim* | X | 103000769-103024890(-1) |
| *Ropn1l* | 15 | 31441357-31453883(-1) |
| *Ror2* | 13 | 53263348-53440160(-1) |
| *Runx2* | 17 | 44806874-45125684(-1) |
| *Scaf11* | 15 | 96309580-96358724(-1) |
| *Scart2* | 7 | 139827197-139880649(1) |
| *Scn3b* | 9 | 40180513-40202914(1) |
| *Sec24b* | 3 | 129776408-129855202(-1) |
| *Sel1l* | 12 | 91772817-91815931(-1) |
| *Selenok* | 14 | 29690265-29697619(1) |
| *Setd3* | 12 | 108072690-108145573(-1) |
| *Setd5* | 6 | 113054326-113130396(1) |
| *Sfr1* | 19 | 47720121-47724027(1) |
| *Sh2d5* | 4 | 137977714-137988643(1) |
| *Shroom3* | 5 | 92831294-93113177(1) |
| *Sik3* | 9 | 45924118-46135492(1) |
| *Sirt3* | 7 | 140443579-140462222(-1) |
| *Skida1* | 2 | 18045487-18053862(-1) |
| *Slc20a2* | 8 | 22966804-23059628(1) |
| *Slc25a1* | 16 | 17743087-17746083(-1) |
| *Slc25a30* | 14 | 75997557-76024477(-1) |
| *Slc25a4* | 8 | 46659834-46664321(-1) |
| *Slc26a2* | 18 | 61325991-61344684(-1) |
| *Slc29a1* | 17 | 45896126-45910532(-1) |
| *Slc29a3* | 10 | 60547851-60588573(-1) |
| *Slc30a9* | 5 | 67464298-67515786(1) |
| *Slc35d1* | 4 | 103027846-103072361(-1) |
| *Slc6a5* | 7 | 49559894-49613604(1) |
| *Slmap* | 14 | 26134323-26256086(-1) |
| *Snx3* | 10 | 42378026-42411377(1) |
| *Spopl* | 2 | 23396232-23462118(-1) |
| *Srd5a3* | 5 | 76288118-76303351(1) |
| *Supt5* | 7 | 28014316-28038171(-1) |
| *Svep1* | 4 | 58042442-58206859(-1) |
| *Tbx20* | 9 | 24629434-24685599(-1) |
| *Tfap4* | 16 | 4362525-4377718(-1) |
| *Tfec* | 6 | 16833372-16898440(-1) |
| *Tgds* | 14 | 118349323-118370167(-1) |
| *Tm9sf4* | 2 | 153003223-153052386(1) |
| *Tmc6* | 11 | 117656814-117673024(-1) |
| *Tmem132a* | 19 | 10835186-10847304(-1) |
| *Tmem248* | 5 | 130245922-130272606(1) |
| *Tmem70* | 1 | 16735431-16748499(1) |
| *Tpte* | 8 | 22773457-22861434(1) |

| | | |
|---|---|---|
| *Traf3ip1* | 1 | 91422369-91457029(1) |
| *Tram2* | 1 | 21066523-21149453(-1) |
| *Trim61* | 8 | 65465639-65471175(-1) |
| *Trip13* | 13 | 74059466-74085903(-1) |
| *Ttc28* | 5 | 111027669-111437646(1) |
| *Twist1* | 12 | 34007670-34009828(1) |
| *Ube2g1* | 11 | 72498109-72577307(1) |
| *Uchl1* | 5 | 66833434-66844577(1) |
| *Ushbp1* | 8 | 71836916-71848446(-1) |
| *Vangl2* | 1 | 171828527-171856011(-1) |
| *Vcan* | 13 | 89803431-89890628(-1) |
| *Vgll3* | 16 | 65612143-65663254(1) |
| *Vps53* | 11 | 75937052-76070473(-1) |
| *Vstm2a* | 11 | 16207724-16377310(1) |
| *Wac* | 18 | 7868832-7929028(1) |
| *Wdr37* | 13 | 8853004-8921945(-1) |
| *Wdtc1* | 4 | 133019770-133080792(-1) |
| *Xaf1* | 11 | 72192455-72204559(1) |
| *Xbp1* | 11 | 5470659-5475893(1) |
| *Xndc1* | 7 | 101714718-101732972(1) |
| *Zdhhc20* | 14 | 58070160-58127733(-1) |
| *Zfhx2* | 14 | 55297719-55329781(-1) |
| *Zmym2* | 14 | 57124110-57200158(1) |
| *Zscan2* | 7 | 80510668-80526285(1) |

Supplemental table 2. The 25 skeletal parameters can be assigned to six groups based on phenotype (see main text for more detail).

| **Somitogenesis** | **Spine shape** | **Tail morphology** | **Vertebral form** | **Vertebral number** | **Vertebral processes** |
|---|---|---|---|---|---|
| Alg10b | Kcnv2 | Col11a1 | Abca4 | Aff4 | Duoxa2 |
| Ankmy2 | Ropn1l | Dis3l2 | Arpc1b | Cacna1c | Fbn2 |
| Atp1b1 | Sik3 | Fgf3 | Bptf | Cfd | Brd2 |
| Atp1b3 | Slc20a2 | Pcgf3 | Cbx5 | Chrna2 | Dcbld2 |
| Ctbp2 | Tpte | Shroom3 | Dnase1l2 | Cx3cr1 | Gal3st1 |
| Eng | Tram2 | Slc26a2 | Fbn2 | Cysltr2 | Glg1 |
| Flnc | Wdr37 | Slc35d1 | Fbrsl1 | Eng | Kera |
| Glmn | Mbd1 | Tmem132a | Fggy | Helq | Runx2 |
| Isyna1 | Ccnd2 | Cant1 | Gm13547 | Lox | Selenok |
| Krit1 | Dnase1l2 | Chd1l | Gna11 | Mllt10 | Tram2 |
| Nog | H2-Eb1 | Cyb561 | Gramd2a | Nog | Twist1 |
| Nr6a1 | Nisch | Dnase1l2 | Gyg | Parp4 | Wdtc1 |
| Opa1 | Pabpc4 | Fgf7 | Il12a | Pde5a | Zfhx2 |
| Pdhb | Selenok | Hoxc12 | Lyplal1 | Slc29a3 | Aff3 |
| Pigq | Uchl1 | Htr1f | Os9 | Trim61 | Arpc1b |
| Rad9a | Vgll3 | Ifi27l2a | Pdss2 | Vstm2a | Cbx5 |
| Sfr1 | | Micu1 | Ppp1r42 | Xaf1 | Dnase1l2 |
| Slc30a9 | | Mir320 | Scn3b | Pcgf2 | Pals2 |
| Tbx20 | | Nono | Sel1l | R3hdm1 | Rlim |
| Tgds | | Npr2 | Setd3 | Rbm22 | Slc25a4 |
| Tmem70 | | Asphd1 | Setd5 | Slc6a5 | Ube2g1 |
| Traf3ip1 | | Barx2 | Sirt3 | Tfap4 | Wac |
| Vcan | | Capza1 | Slc29a1 | | Wdr37 |
| Zmym2 | | Fbn2 | Tmc6 | | Xndc1 |
| | | Fbrsl1 | Tmem248 | | C9orf72 |
| | | Fbxo22 | Ttc28 | | 4932438H23Rik |
| | | Mmp11 | Ube2g1 | | Gldc |
| | | Pfn4 | Ushbp1 | | Pld5 |
| | | Pramel17 | Wdr37 | | Ralb |
| | | Pstpip2 | Zscan2 | | Sh2d5 |
| | | Scart2 | Barx2 | | Skida1 |
| | | Tfec | Colec10 | | Slc25a30 |
| | | Trip13 | Cpt2 | | Spopl |
| | | Cdcp3 | Cyp27b1 | | Vps53 |
| | | Zdhhc20 | Dbn1 | | Xbp1 |
| | | Coq6 | Dcdc2c | | |
| | | Glmn | Duoxa2 | | |
| | | Ipo9 | Hip1 | | |
| | | Isyna1 | Il2ra | | |
| | | Med11 | Klf7 | | |
| | | Ndufb5 | Lrrk1 | | |
| | | Nr6a1 | Mtch2 | | |
| | | Pigq | Notch1 | | |
| | | Ppp1r35 | Plekhm1 | | |
| | | Slc30a9 | Pstpip2 | | |
| | | Srd5a3 | Runx2 | | |
| | | Ambra1 | Sik3 | | |

| | | | |
|---|---|---|---|
| | | Gdf11 | Twist1 |
| | | Ino80c | 1110059G10Rik |
| | | Ndel1 | Aff3 |
| | | Pcsk5 | Cog6 |
| | | Psen1 | Dph6 |
| | | Slc25a1 | Hbs1l |
| | | Aebp1 | Jmjd1c |
| | | Dpf2 | Kdm7a |
| | | Fgfr1op2 | L3mbtl2 |
| | | Fuz | Mau2 |
| | | Lum | Mbd1 |
| | | Pfdn5 | Pitx1 |
| | | Ror2 | Scaf11 |
| | | Sec24b | Supt5 |
| | | Snx3 | Tm9sf4 |
| | | Vangl2 | |
| | | Adamts1 | |
| | | Celsr1 | |
| | | Cgn | |
| | | Gramd1b | |
| | | Nemf | |
| | | Pfdn1 | |
| | | Rbm45 | |
| | | Rexo1 | |
| | | Slmap | |
| | | Svep1 | |

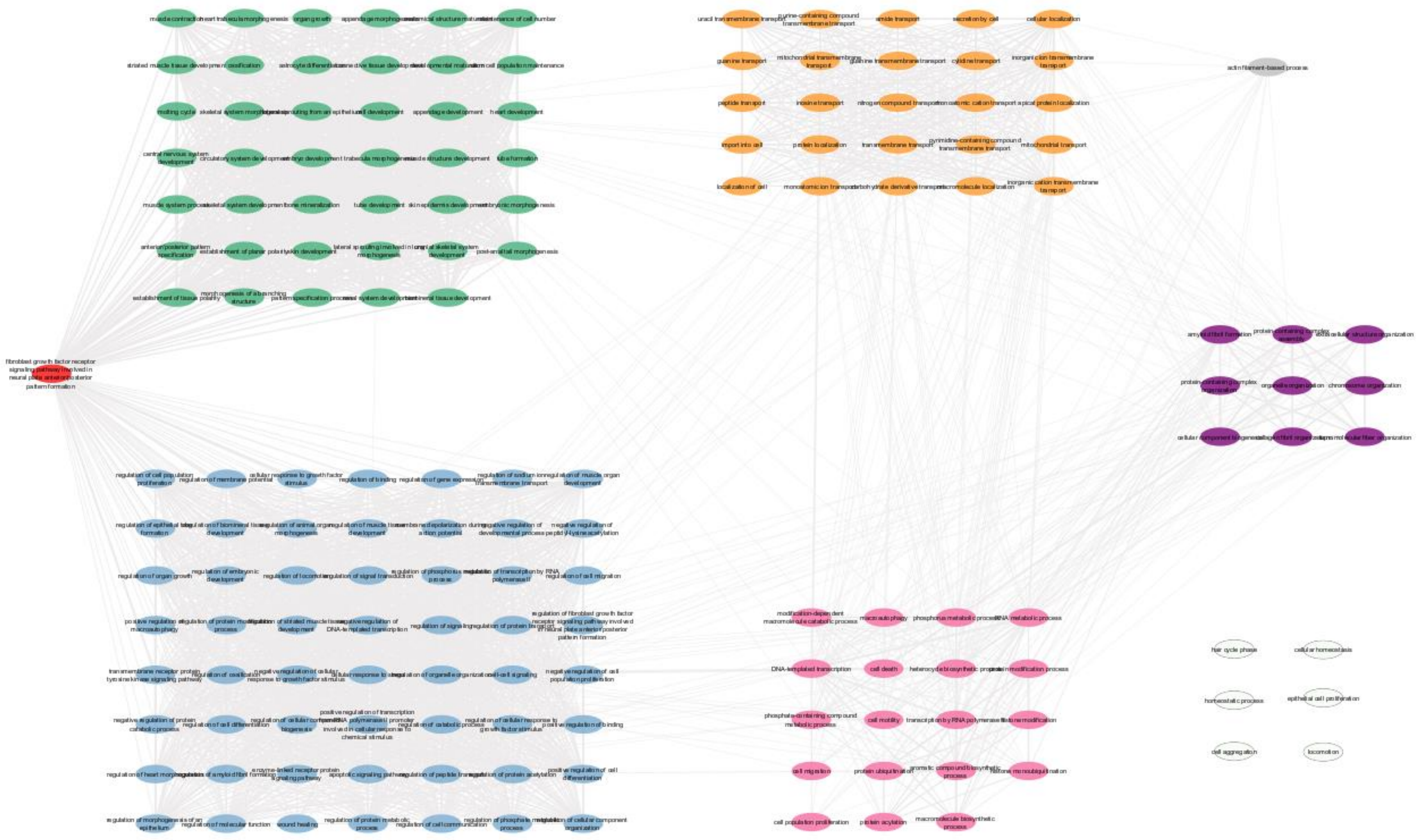

Supplemental figure S1. ReviGO clustering of Gene Ontology (GO) terms, based on DAVID functional annotation (GOTERM_BP_FAT, p<0.05). Development (green); metabolism (pink); transport (orange); regulation (blue); molecular organisation (purple).

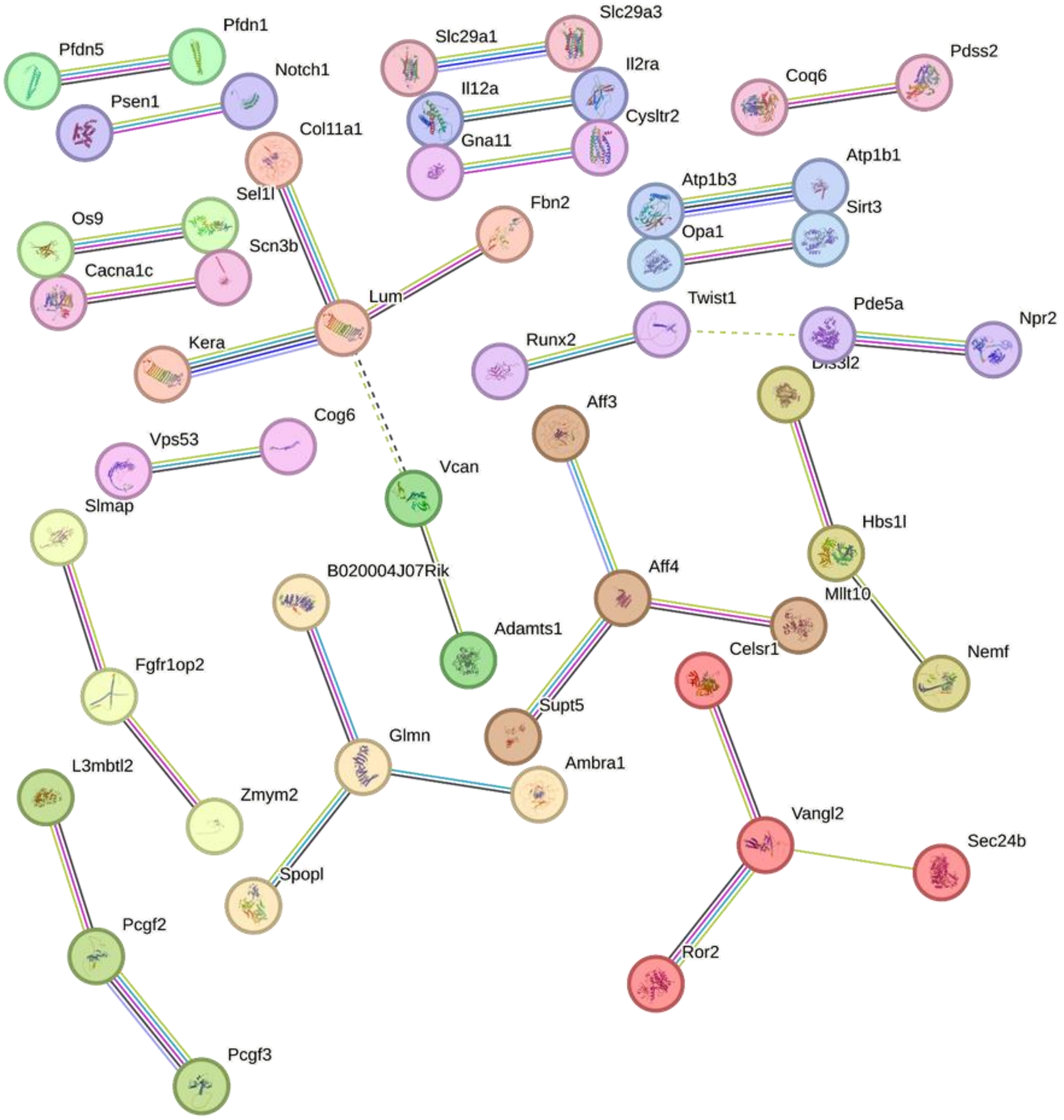

Supplemental figure S2. Analysis of protein:protein interactions within the 204 IMPC genes with a vertebral phenotype using the STRING biological database, with Markov Cluster Algorithm (MCL), showing the interactions with high (0.70–0.89) confidence.

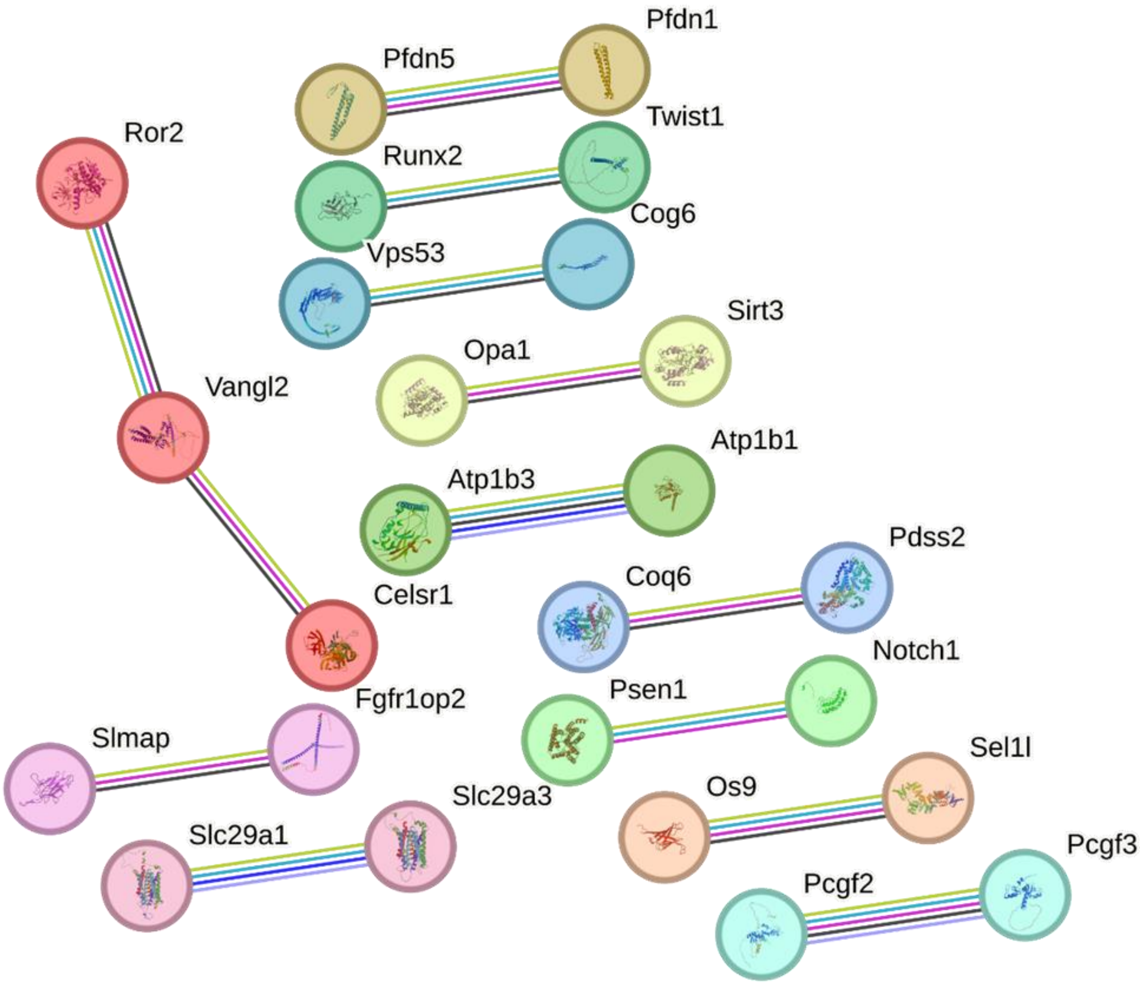

Supplemental figure S3. Analysis of protein:protein interactions within the 204 IMPC genes with a vertebral phenotype using the STRING biological database, with Markov Cluster Algorithm (MCL), showing the interactions with highest (0.90–1.0) confidence.
